# SACNet: A Multiscale Diffeomorphic Convolutional Registration Network with Prior Neuroanatomical Constraints for Flexible Susceptibility Artifact Correction in Echo Planar Imaging

**DOI:** 10.1101/2023.09.15.557874

**Authors:** Zilong Zeng, Jiaying Zhang, Xinyuan Liang, Lianglong Sun, Yihe Zhang, Weiwei Men, Yanpei Wang, Rui Chen, Haibo Zhang, Shuping Tan, Jia-Hong Gao, Shaozheng Qin, Qiqi Tong, Hongjian He, Sha Tao, Qi Dong, Yong He, Tengda Zhao

**Author notes:** **Corresponding author:** Tengda Zhao, Ph.D.

## Abstract

Susceptibility artifacts (SAs), inevitable in brain diffusion MR (dMRI) scans acquired using single-shot echo planar imaging (EPI), severely compromise the accurate detection of human brain structure. Existing SA correction (SAC) methods offer inadequate correction quality and limited applicability across diverse datasets with varied acquisition protocols. To address these challenges, we proposed SACNet, a SAC framework based on unsupervised registration convolutional networks, featuring: i) a novel diffeomorphism regularization function to avoid unnatural SAC warps, modified from a potential well function; ii) an integration with prior neuroanatomical constraints and coarse-to-fine processing strategy to enables multi-scale geometric and intensity recoveries in severe distorted areas; iii) a unified registration framework that incorporates multiple phase-encoding (PE) EPI images and structural images, ensuring compatibility with both single- and inverse-PE protocols, with or without field maps. Utilizing simulated dMRI images and over 2000 brain scans from neonatal, child, adult and traveling participants, our method consistently demonstrates state-of-the-art correction performance. Notably, SACNet effectively reduces SAs-related multicenter effects compared to existing methods. We have developed user-friendly tools using containerization techniques, hope to facilitate SAC correction quality across extensive neuroimaging studies.

## 1. Introduction

Diffusion magnetic resonance imaging (dMRI) provides a unique opportunity to noninvasively probe human brain white matter (WM) in vivo (Hagmann, 2005; Lerch et al., 2017; Sporns et al., 2005), which is crucial for modern neuroscience studies. To improve imaging speed and achieve high image resolution (Glasser et al., 2013; Littlejohns et al., 2020; Somerville et al., 2018), dMRI scans usually employ the single-shot echo planar imaging (EPI) sequence (Biswal et al., 1995; Turner et al., 1990; Warach et al., 1995), which has become the predominant choice for neuroimaging studies, especially in recent large-scale cohort projects. However, this approach is substantially affected by susceptibility artifacts (SAs), resulting in severe geometric and intensity distortions (Andersson et al., 2003; Jezzard and Balaban, 1995), which largely confound accurate measurements of brain WM from voxel-wise to connectome level brain studies (Tax et al., 2022). Evidence from multicenter datasets showed that SAs lead to the largest inconsistency in brain connectivity measurements across scan centers (Yamashita et al., 2019). Developing a high-quality susceptibility artifact correction (SAC) approach for diverse datasets is an ongoing task for brain neuroimaging studies.

Conventional approaches for solving the SAC problem generally employ two frameworks: the single-phase encoding (single-PE) based method and the inverse phase encoding (inverse-PE) based method. Both of them depend on specific EPI protocol designs. The single-PE approach usually requires an additional scan of raw magnetic field inhomogeneity, derived from the dual-echo field map sequence (Jezzard and Balaban, 1995; Reber et al., 1998). SAs are corrected by translating the dual-echo field map into local voxel shifts. The inverse-PE approach relies on two PE-opposite EPI scans to capture the complementary signal along inversed distortion directions (Andersson et al., 2003; Bowtell et al., 1994; Hédouin et al., 2017; Holland et al., 2010; Irfanoglu et al., 2015; Ruthotto et al., 2012). SAs are corrected by finding an ideal “middle” estimation between two inversed distorted images through iterative registrations. The most recognized method of the inverse-PE approach is “Topup” in FSL software (Andersson et al., 2003), which presents a least-squares estimation of opposing undistorted images and usually outperforms the field map framework. However, these methods suffer from common drawbacks, including the limited performance which caused by cumulative errors in iterative registration, the narrow applicability restricted to specific EPI sequence designs, and the rather long computation time.

Recently, new SAC approaches utilizing convolutional neural networks (CNNs) have emerged and enabled faster and superior SAC performance than traditional approaches in various EPI protocols. Methodologically, they can be mainly classified into two categories: synthetic models (Hu et al., 2020; Ye et al., 2023) and registration models (Duong et al., 2020b; Qiao and Shi, 2021; Zahneisen et al., 2020). The synthetic models collect additional distortion-free adult brain images (specialized MRI protocols, such as point-spread-function (PSF)–encoded EPI) as supervised labels (Hu et al., 2020; Ye et al., 2023). Such methods permit SAC on single-PE images without requiring a field map. But they are inherently limited by the supervised training process, which may lead to inadequate performance for out-of-sample brain images with pronounced variations, such as neonatal or multicenter brain scans. Existing registration-based models are mainly designed for inverse-PE images (Duong et al., 2020b; Qiao and Shi, 2021; Zahneisen et al., 2020). Such methods enables learning common spatial mappings to the undistorted brain images via neural network training process thus could avoid the iterative registration process (Balakrishnan et al., 2019). The self-supervised learning process also brings high generalization ability, which is critical for robust SAC performance on various EPI protocols. However, several limitations still exist for current registration-based CNN models: 1) a compatible framework that can handle both single- and inverse-PE type datasets, with or without field-map acquisitions, is lacking, particularly for multi-cohort datasets containing distinct protocol designs; 2) failing to ensure diffeomorphic transformations leads to artificial warps during distortion correction; 3) prior neuroanatomical information from structural MR images is underestimated; and 4) the single-resolution strategy hampers model convergence.

To fill these gaps, we proposed SACNet, an unsupervised CNN registration method for SAs correction with the following innovations:

i. We established a mathematical framework to flexibly tackle the SAC problem under both inverse-PE and single-PE EPI protocols by integrating multiple PE images and structural images within a unified registration pipeline.
ii. We proposed a diffeomorphism preservation regularization function by modifying the Woods-Saxon potential function to restrict the generated deformation fields within a diffeomorphic solver space.
iii. We designed an intensity-irrelevant loss function that is suitable for both T1w and T2w brain images to introduce anatomical priors for recovering cortical details in severely distorted region and ensure usability for dataset without fieldmap acquisition.
iv. We designed coarse-to-fine (CTF) training and inference strategy to accelerate the learning process, leading to satisfactory model convergence.

By employing simulated dMRI images as well as nearly 2000 brain dMRI images covering neonatal, child, adult and traveling participants across multiple scan centers, our SACNet consistently outperforms both conventional and deep-learning based methods in all datasets with significantly improved correction performance, reduced multicenter effects, and low computational costs. We integrated our model into a public preprocessing pipeline at https://github.com/RicardoZiTseng/SACNet. This paper is organized as follows. In Section 2, we describe the detailed network design. In Section 3, we introduce the datasets, evaluation metrics and baseline models. In Section 4, we present experimental results for all datasets. In Section 5, we discuss the results.

## 2. Methods

Firstly, we present an overview of the SACNet framework (Section 2.1, Fig. 1). Then, we detailly describe the differentiable EPI warp module (Section 2.2), the mathematical optimization functions (Section 2.3), the formulated optimization model and its variants for different PE protocols (Section 2.4), and the CTF training and inference approach (Section 2.5). Finally, we introduce how to integrate SACNet into common public dMRI preprocessing pipelines (Section 2.6).

**Fig. 1.**
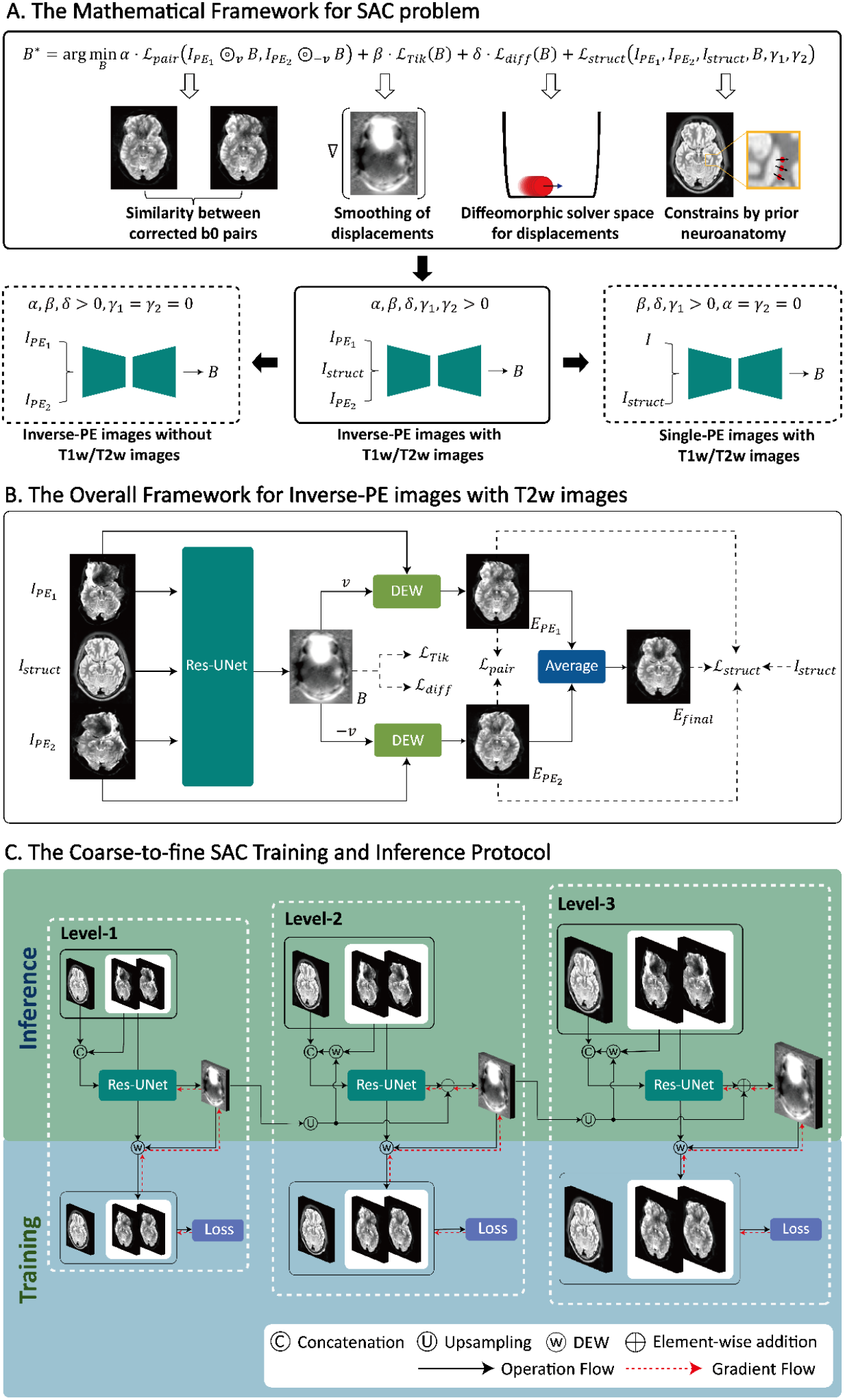
The proposed mathematical framework and an implementation flowchart of SACNet. (A) The first row presents the integrated mathematical optimization function for solving the SAC problem, including a pairwise dissimilarity loss function ℒ_*pair*_, a Tikhonov regularization function ℒ_*Tik*_, a diffeomorphism preservation regularization function ℒ_*diff*_, and a prior neuroanatomical information loss ℒ_*struct*_. The second row illustrates that our proposed framework can be decomposed into three models, allowing SACNet to be compatible with diverse PE protocols by strategically adjusting the hyperparameters. (B) The example implementation framework of SACNet with inverse-PE b0 images and T2w images as inputs is shown. All input images were sent to Res-UNet to map the inhomogeneity field *B* for correcting SAs. The solid line represents the data flow in the network, and the dashed line represents the participation in the loss function calculation. (C) The implementations of CTF SAC training and inference protocols for the model presented in (B). We used a series of identical networks to simulate the SAC process in the multiresolution schema. The blue part illustrates the optimization of network parameters during the training stage, and the green part illustrates the data flow during the inference stage.

### 2.1 Overview

The mathematical framework and a representative flowchart of SACNet are illustrated in Fig. 1A and Fig. 1B. To flexibly solve the SAC problem for various imaging protocols, we designed a unified registration pipeline compatible for both inversed-PE EPI registration and single-PE EPI to T1w/T2w structural image registration. To obtain natural and accurate SAC warps, we proposed an integrated optimization function incorporates four loss functions: a pairwise dissimilarity loss function ℒ_*pair*_, for cases with an available inverse-PE b0 (b-value=0 s/mm^2^) image pair, a Tikhonov regularization function ℒ_*Tik*_, for generating smooth inhomogeneity fields, a diffeomorphism preservation regularization function, ℒ_*diff*_, for guaranteeing diffeomorphic inhomogeneity fields, and a prior neuroanatomical information loss, ℒ_*struct*_, for incorporating prior neuroanatomical information. ℒ_*struct*_ also served as registration target when only single-PE image is available. Overall, this mathematical framework can be decomposed into three models (Fig. 1A, bottom panel), enabling SACNet to be compatible with diverse PE protocols by strategically adjusting hyperparameters.

To illustrate this framework in detail, here we took the full model (for inverse-PE images with T2w images as the prior neuroanatomical information) as an example (Fig. 1B). Specifically, we employed a Res-UNet to model the mapping from 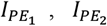 and *I*_*struct*_ to *B*: 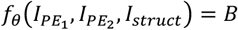, in which 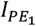 and 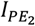 are the uncorrected image pair along the inverse-PE directions, *I*_*struct*_ is the structural image rigidly registered to 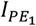 and 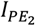, *B* is the generated inhomogeneity field needed to remove SAs, and *θ* represents the network parameters. The Res-UNet layout was consisted of an encoder-decoder design with skip connections links and pre-activation (PreAct) block (SI-4). A differentiable EPI warp (DEW) module was designed to implement both geometric and intensity corrections and generate the corrected images along the two PE directions, 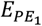 and 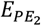. Finally, we combined 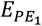 and 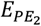 based on the average to generate the final corrected image *E*_*final*_. For faster and better training convergence, we designed CTF training and inference protocols, as shown in Fig. 1C.

### 2.2 The differentiable EPI warp module

To enable simultaneously implementation of geometry and intensity corrections, we constructed a differentiable EPI warp module by designing image calculations process based on previous physical imaging models. Specifically, for image *I*, we first computed the voxel location ***p*′** **= *p* +** *B*(***p***)***v*** for each voxel ***p*** in image *I*. By assuming that SAs only occur along the PE direction, we linearly interpolated the values for the left-right neighboring voxels along the PE direction ***v***:

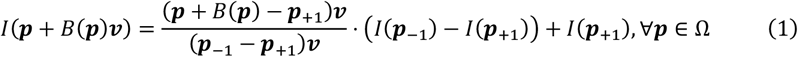

where ***p***_−1_ and ***p***_**+**1_ are the previous and next neighbor voxels of ***p*** along the PE direction ***v***. Then, we multiplied Eq. (1) by the Jacobian determinant of ***B*** to redistribute the intensity as follows:

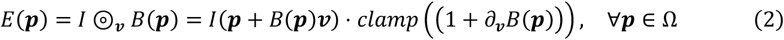

where (1 **+** ∂_***v***_*B*(***p***)) in Eq. (2) is the Jacobian determinant of the transformation ***p*** → ***p* +** *B*(***p***)***v*** (see detailed derivation in SI-1), and *clamp*(*x*) **=** *max*(*x*, 0) is used to prevent multiplication with a negative value.

### 2.3 Optimization function construction

For images with inverse-PE designs, the distorted image pair 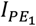 and 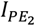 is inversely affected by the same inhomogeneity field *B* along the opposite directions ***v*** and −***v*** (Holland et al., 2010; Ruthotto et al., 2012); thus, the corrected images 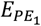 and 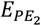 were calculated as follows according to Eq. (2):

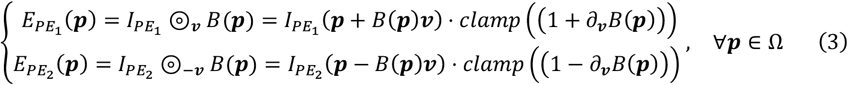

Theoretically, we can find one solution *B*^∗^ that leads to identical 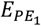 and 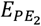; thus, the optimization problem can be formulated as:

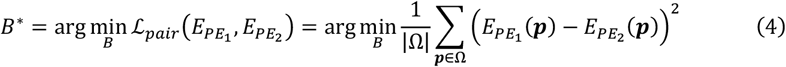

where ℒ_*pair*_ adjusts the pairwise dissimilarity between the estimated 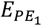 and 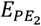. Notably, all image volumes are defined over a 3D spatial domain Ω ⊂ ℝ^3^, and |Ω| represents the number of elements in Ω.

However, previous studies have noted that seeking *B*^∗^ by optimizing Eq. (4) generally leads to an ill-posed problem and is inadequate for severe distortions (Balakrishnan et al., 2019; Duong et al., 2020a; Ruthotto et al., 2012). We introduced two regularization functions (ℒ_*Tik*_and ℒ_*diff*_) and one structural constrain function (ℒ_*struct*_) to constrain the solver space of *B*:

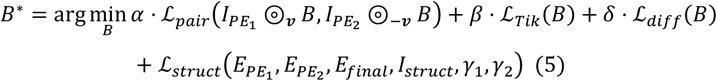

where *α*, *β*, *δ*, *γ*_1_ and *γ*_2_ are hyperparameters used to determine the contribution of each component in Eq. (5). In addition, ℒ_*Tik*_, ℒ_*diff*_, and ℒ_*struct*_ denote the Tikhonov regularization function, diffeomorphism preservation regularization function and prior neuroanatomical information loss function, respectively, which are defined in the subsequent subsections.

#### 2.3.1 Tikhonov regularization

ℒ_*Tik*_(*B*) was used as a prior constraint on the smoothness of field *B* using a Tikhonov regularizer based on the spatial gradient of *B*:

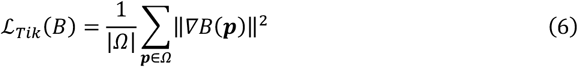

Following the implementations in (Balakrishnan et al., 2019), for 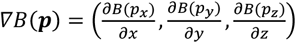, we approximated 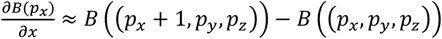, and we used similar approximations for 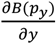 and 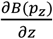.

#### 2.3.2 Diffeomorphism preservation regularization

To guarantee the diffeomorphism of the inhomogeneity field, we proposed a diffeomorphism preservation regularization function by modifying a potential well function. Specifically, in terms of intensity, we expected that the signals of the voxels at the same position in *E*_1_ and *E*_2_ would both be positive, which requires the following:

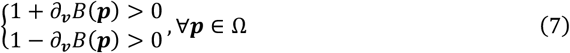

This is equivalent to:

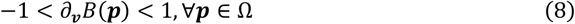

In terms of the geometry, we expected the relative positions of adjacent voxels to remain the same before and after resampling, which guarantees no folding areas during the transformation, as shown in Fig. 2A. To this end, the new spatial positions should follow:

**Fig. 2.**
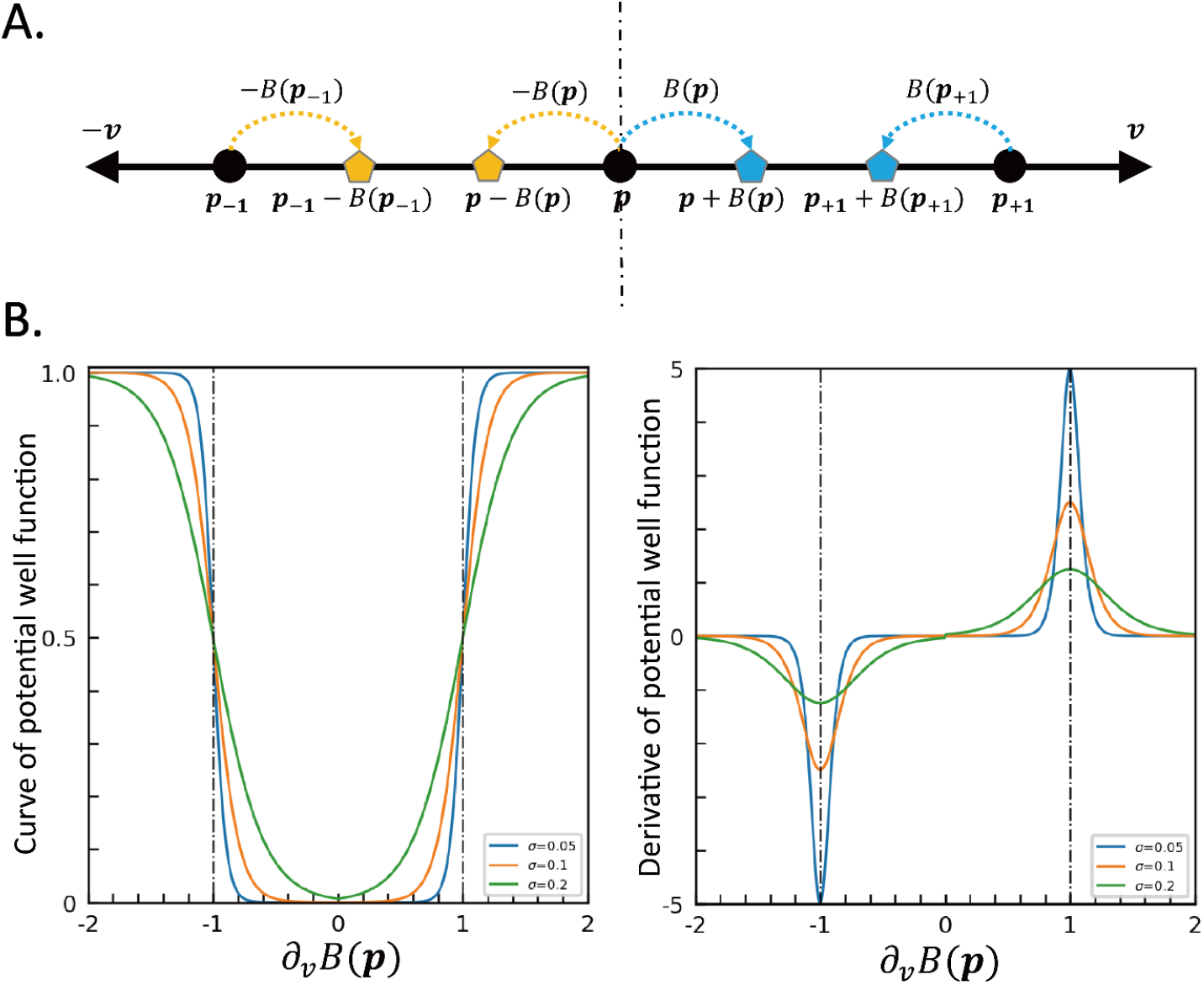
Mathematical framework of the diffeomorphism preservation regularization. (A) Illustration of spatial folding at location *x* along opposite directions ***v*** and −***v***. (B) The left part shows the function value of *ϕ*(*x*) with different hyperparameters σ and the value in terms of ∂_***v***_*B*(***p***). The right part shows the derivative of *ϕ*(*x*) with respect to ∂_***v***_*B*(***p***).

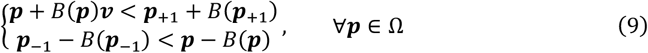

We can also obtain Eq. (8) from Eq. (9).

To ensure that the generated field *B* satisfies the constraint function shown in Eq. (8), we expected that when ∂_*v*_*B*(*p*) approached -1 or 1, the loss function increased substantially, and when ∂_*v*_*B*(*p*) remained between -1 and 1, the loss function remained small. To this end, we designed the diffeomorphism preservation function (DPF) as follows:

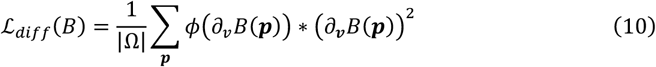

where *ϕ*(·) is the potential well function modified from the Woods-Saxon potential function widely used in nuclear physics (Erkol and Demiralp, 2007):

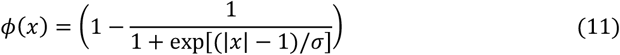

where σ is a customized parameter. Fig. 2B shows the curves of *ϕ*(*x*) (the left subgraph) in terms of ∂_***v***_*B*(***p***) and the derivative (the right subgraph) with respect to ∂_***v***_*B*(***p***). The figure shows that the value of *ϕ*(*x*) increases substantially as |∂_***v***_*B*(***p***)| → 1, which suggests that *ϕ*(*x*) can sensitively suppress the voxels that do not obey the constraint defined in Eq. (8), thereby constraining the inhomogeneity field *B* to a diffeomorphic space. Notably, (∂_***v***_*B*(***p***))^2^ is multiplied with *ϕ*(∂_***v***_*B*(***p***)) to prevent the gradient from vanishing when ∂_***v***_*B*(***p***) is larger than 1

or smaller than -1. We present the proof of the existence of a diffeomorphic inhomogeneity field calculated by SACNet in SI-2.

#### 2.3.3 Prior neuroanatomical information loss

Image noise caused by SAs hinders strict alignment of the b0 image pair, resulting in inaccurate estimation of the inhomogeneity field in severely distorted areas. To address this issue, we proposed a prior neuroanatomical information loss ℒ_*struct*_ to incorporate accurate prior neuroanatomical information. This approach has two main benefits. First, this loss regularizes the inhomogeneity field while preserving intricate neuroanatomical morphological details. Second, it provides an additional registration target when the b0 image pair is not available.

ℒ_*struct*_ consists of two parts: the overall shape structural similarity loss ℒ_*str*−*overall*_ and the pairwise structural similarity loss ℒ_*str*−*pair*_ . Conceptually, ℒ_*str*−*overall*_ ensures that the final corrected image *E*_*final*_ is similar to the provided structural image *I*_*struct*_, while ℒ_*str*−*pair*_ ensures that the corrected images along each PE direction 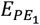 and 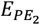 are similar to *I*_*struct*_. Specifically, the proposed neuroanatomy prior loss is formulated as:

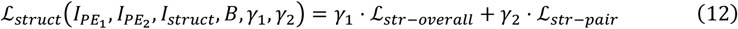

With

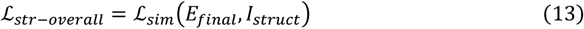

And

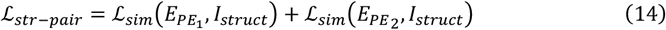

where *γ*_1_ and *γ*_2_ are two user-defined hyperparameters, and ℒ_*sim*_ in Eq. (13) and Eq. (14) represents a similarity metric.

We anticipated that SACNet would not be limited to the MR modality of structural inputs. Therefore, instead of relying on absolute intensity-relevant similarity metrics, such as the mean square error (MSE) and local cross-correlation (LCC), we employed a gradient-based similarity metric, namely, the normalized gradient field (NGF), as ℒ_*sim*_. The NGF determines the geometric resemblance between any points in an image by computing local gradients; thus, this metric is independent of the absolute image intensity (Haber and Modersitzki, 2007). Let ∇*X*_***p***_ be the intensity change gradient at point ***p*** ∈ Ω in image *X* and ∈ be a user-defined parameter that prevents divide-by-zero errors. Then, the NGF measure at any point ***p*** in image *X* can be defined as:

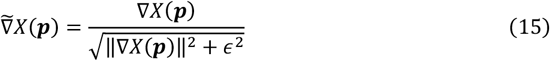

The difference between two images *X* and *Y* can be measured by calculating the angles between the NGF vectors at all points in the image domain, which can be formulated as follows:

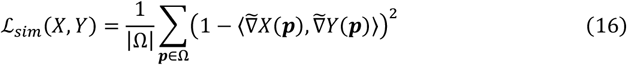

where ⟨·,·⟩ denotes the inner dot-product operation. The value of ℒ_*sim*_(*X, Y*) is positive, and the smaller the value of ℒ_sim_(*X, Y*) is, the more similar the two images are.

### 2.4 The formulated optimization model and its variants

To handle the different imaging protocols in various existing neuroimaging datasets, the optimization model formulated in Eq. (5) can be transformed into two different forms, as shown in Fig. 1A: a) When no structural images are available (image set 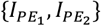), the model can be transformed to use Eq. (17) by setting *γ*_1_ and *γ*_2_ to 0, as illustrated in the first column in the second row of Fig. 1A:

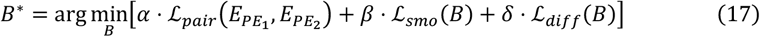

b) When only single-PE images are available (only one single-direction distorted image and one structural image, with the image set {*I, I*_*struct*_}), the model can be transformed to use Eq. (18) by setting *α* and *γ*_2_ to 0, as illustrated in the third column in the second row of Fig. 1A:

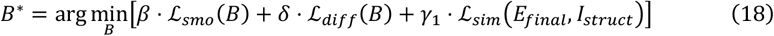

In this situation, *E*_*final*_ **=** *I* ⊚_***v***_ *B* denotes the image corrected based on the distorted image *I* along the single-PE direction ***v*** . In addition, the potential well function in Eq. (11) can be reformulated as:

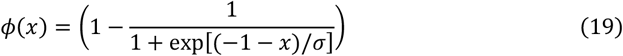

In summary, the overall loss functions of network training for three image sets including 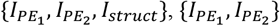 and {*I, I*_*struct*_} are separately shown as follows:

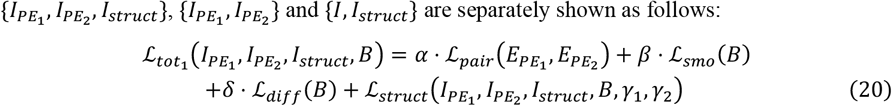

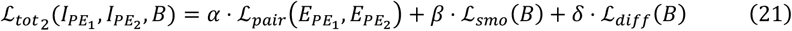

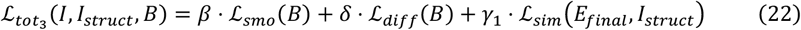

### 2.5 Coarse-to-fine (CTF) SAC training and inference protocols

To improve the training process and prevent falling into local minima, we designed CTF training and inference protocols for SACNet, as illustrated in Fig. 1C. The CTF training protocol aims to train multiple networks at *N*_*s*_ scale levels, with each model estimating the residual inhomogeneity field at the corresponding scale. Specifically, we first trained the network at the coarsest scale level and then progressively trained the networks at each subsequent scale level to solve the SAC problem at finer scale levels. This training procedure was repeated until the model was trained at the finest level. The CTF inference protocol aimed to generate the estimated inhomogeneity field based on the training protocol using multiple trained networks. At each scale level *s*, we downsampled the image set by 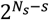 times and upsampled the inhomogeneity field *B*^(*s*−1)^ 2 times. Then, we fed the downsampled image set into the network at the current level to obtain the residual inhomogeneity field Δ*B*^(*s*)^ . The inhomogeneity field at the current level was calculated by summing the upsampled field and the residual field. The pseudocodes for the training and inference protocols are presented in Algorithms 1 and 2, respectively.

#### Algorithm 1. Coarse-to-fine SAC training protocol of SACNet, as depicted in Fig. 1C.

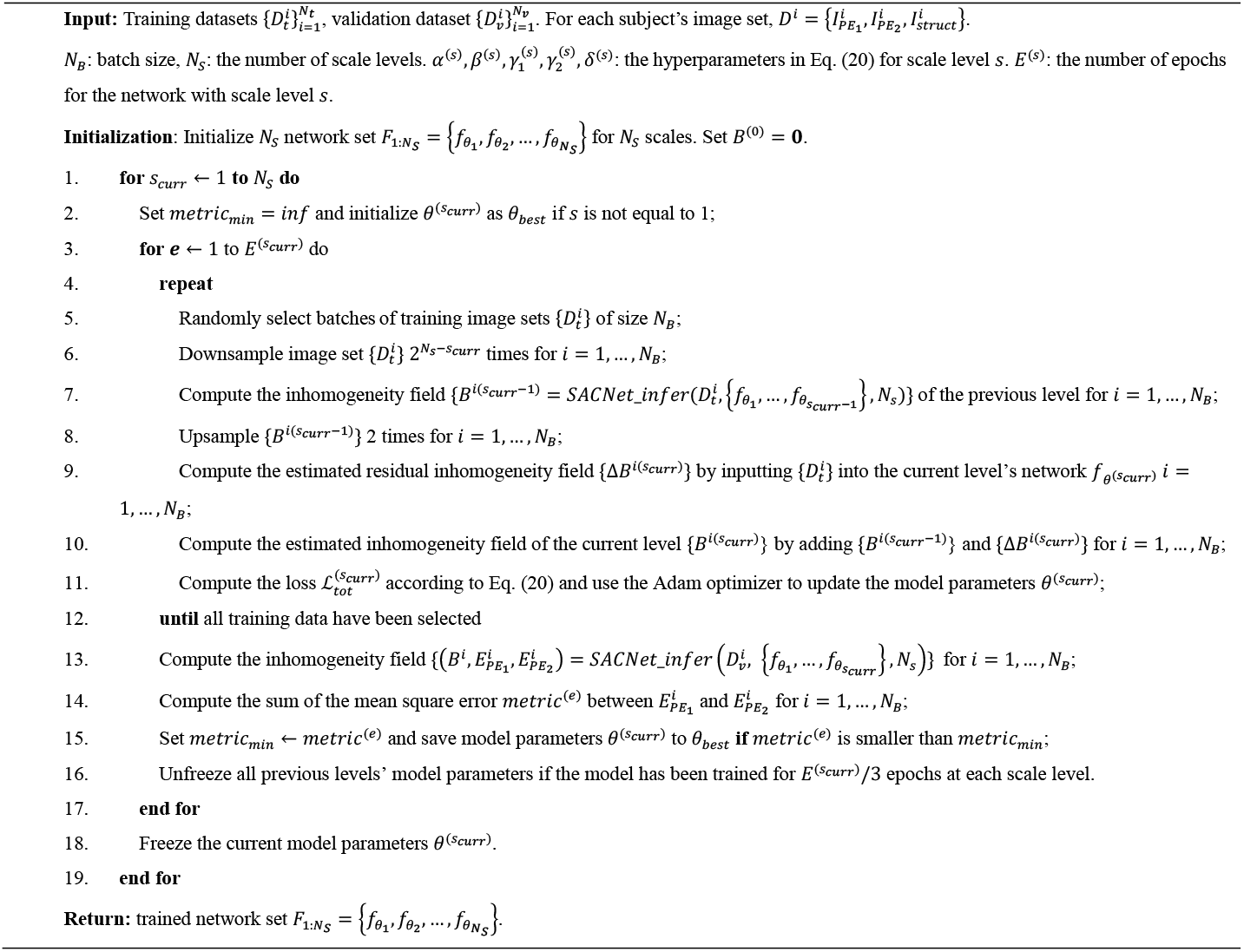

#### Algorithm 2. Coarse-to-fine SAC inference protocol of SACNet, as depicted in Fig. 1C.

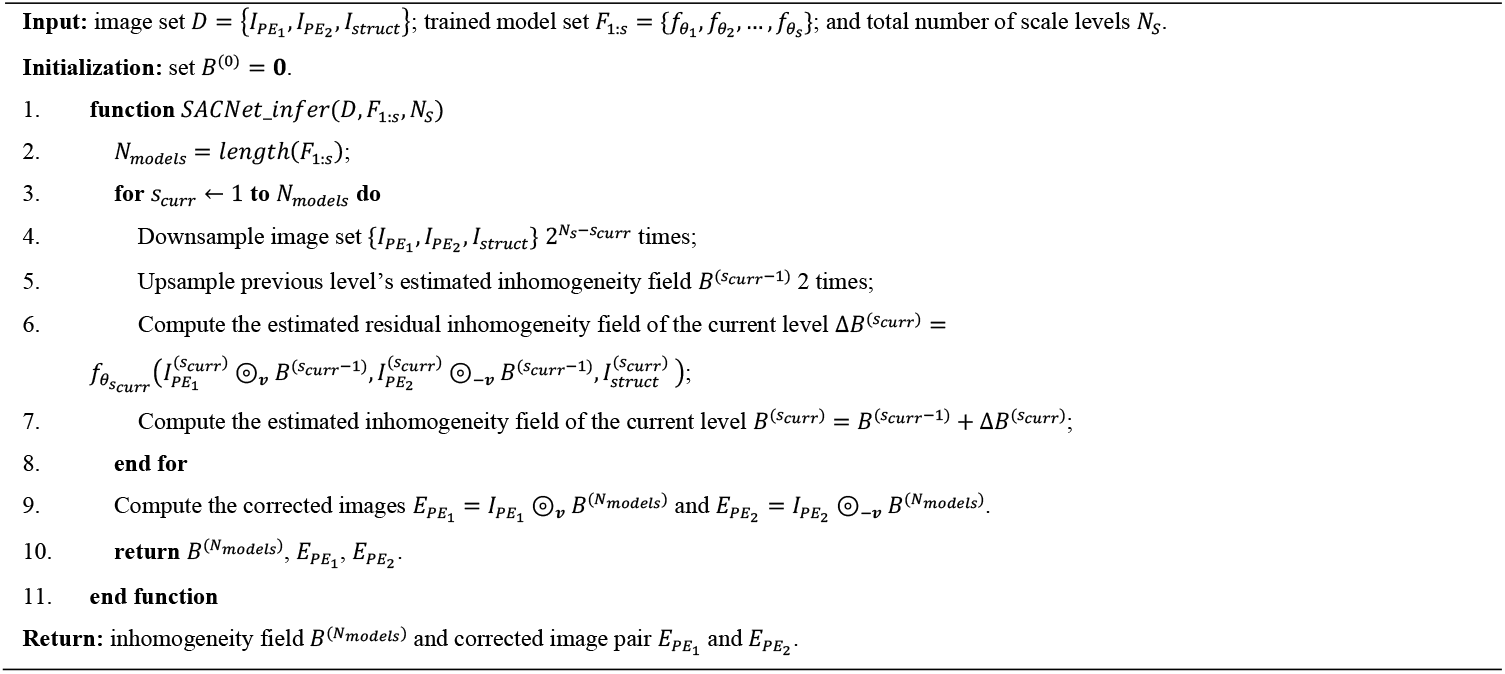

### 2.6 The whole dMRI preprocessing pipeline

We introduced a practical dMRI preprocessing pipeline by integrating SACNet with the popular Eddy tool in FSL. The pipeline started by correcting for motion and eddy current distortions in the dMRI volumes along each PE direction using the FSL Eddy tool. Next, the structural (T1w and T2w) images, as well as all negative and positive PE scans, were rigidly co-registered using the FSL FLIRT tool, with the first b0 image serving as the target. Then, the aligned b0 images were input into the trained model to estimate the inhomogeneity field. Finally, we used the differentiable EPI warp module to remove SAs in all diffusion weighted images (DWIs).

## 3. Experimental settings

### 3.1 Datasets

To comprehensively evaluate the performance of SACNet, we utilized both simulated images and multiple existing neuroimaging datasets spanning various age groups and acquisition protocols:

1. Simulated dMRI images (Simulation): To test our model for dMRI images in an ideal imaging condition, we employed a publicly available dataset with simulated dMRI scans. These images were generated by solving the Bolch and Maxwell equations, ensuring key features that resemble real-world counterparts (Graham et al., 2017). The whole dataset includes a T2w image, a dMRI scan without susceptibility artifact serving as the gold standard, a dMRI scan with susceptibility artifact and a corresponding fieldmap.
2. Inverse-PE dMRI images of adult brain: To test our model for dMRI images with inverse-PE, we randomly selected 380 adult dMRI scans (mean age: 28.8±3.7 years) from the Human Connectome Project (HCP) dataset (Glasser et al., 2013).
3. Inverse-PE dMRI images of developmental brain: To evaluate our model on inverse-PE images of children and neonatal brain, we randomly selected 644 children’s scans (mean age: 13.8±3.8 years) from the Lifespan Human Connectome Project in Development (HCP-D) dataset (Somerville et al., 2018) and 444 neonatal scans (mean gestational age: 39.5±3.3 weeks) from the Developing Human Connectome Project (dHCP) dataset (Makropoulos et al., 2018).
4. Single-PE dMRI images of developmental brain: To test our model on single-PE brain images, we employed brain scans from the Children School Functions and Brain Development Project in China (CBD). We included two independent subsets (CBDP, the Peking University site; CBDH, the Huilongguan Hospital site) totally including 456 children brain scans (mean age: 9.2±1.5 years).
5. Spin-echo (SE) field maps: Current neuroimaging projects, such as the HCP and Baby Connectome Project (BCP), recommend collecting spin-echo EPI (SE-EPI) images with reversed PE directions (SE field maps) to correct SA distortions. Such images could provide good neuroanatomical details and are compatible for both dMRI and fMRI brain images, contributing to their growing popularity. To evaluate the generalization performance of our framework on SE field maps even for fMRI scans. We utilized fMRI SE-EPI field maps from 379 subjects in HCP dataset.
6. The Multicenter dMRI dataset: To assess the performance and reliability of our model across scan sites, we employed 30 brain scans of three healthy traveling subjects (age: two 23 years and one 26 years) acquired at 10 scanning sites from a multicenter public dataset (Tong et al., 2020).

The above datasets were collected in compliance with ethical standards and approved by relevant institutional review boards in each project. All participants or their legal guardians provided informed consent prior to participation. For detailed information about each dataset, see Table 1. Additionally, the preprocessing methods for each dataset are described in SI-3.

**Table 1.**
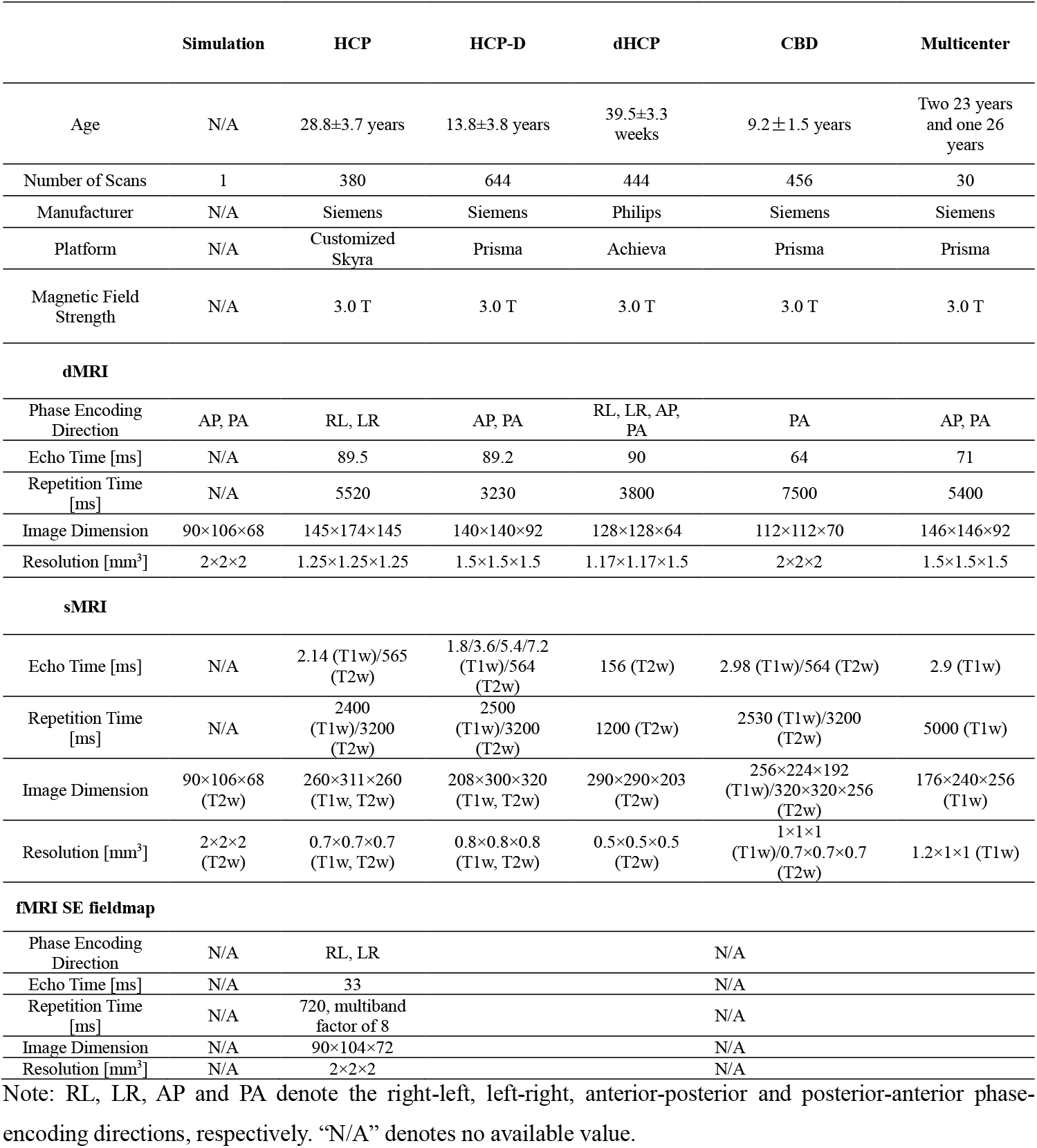
The acquisition parameter details of each dataset.

### 3.2 Evaluation and statistical methods

We adopted multiple evaluation metrics separately tailored to each type of dataset as follows:

#### 1) Simulation dMRI images

We directly calculated the mean square difference between the ground truth and the corrected outputs as the correction error. To give a system evaluation, this procedure is conducted for three types of images including the b0 image, the fractional anisotropy (FA) image and the fieldmap image. Three metrics were generated correspondingly as the b0 error (b0-Err), the FA error (FA-Err) and the fieldmap error (FM-Err) with lower values indicating better performance. To further assess the diffeomorphism of the calculated transformation field, we measured the number of folding voxels (NFV), with lower NFV values indicating better performance.

#### 2) Realistic dMRI dataset

To quantitatively assesses SAC performance for diffusion model fitting, we calculated three metrics based on the FA images including the FA-based mean squared difference between different PE directions (FA-MSD; lower values indicate better performance), FA-based standard deviation across multiple PE directions (FA-STD; lower values indicate better performance), and FA-based structural similarity (FA-SS; local cross-correlation between FA and T1w images with window size of 3; higher values indicate better performance). For the HCP, HCP-D and multicenter datasets, we used the FA-MSD and FA-SS as evaluation indices. For the dHCP dataset, which has four unique PE directions and lacks a sufficient number of structural images, we chose FA-MSD and FA-SS as evaluation indices. For the CBD dataset, which has only one PE direction, we selected the FA-SS as the evaluation index. For all datasets, we also calculated the number of folding voxels (NFV) to assess the extent of diffeomorphism within transformation maps. Of note, to simplify the experiment, we did not utilize the b0 image for the entire evaluation, as it cannot reflect the correction quality in diffusion weighted volumes. However, we calculated the b0-MSD (lower values indicate better performance) and b0-SS (higher values indicate better performance) for the corrected images of HCP dataset for a validation. The results are highly consistent with the FA-based metrics (shown in Supplementary Table S1 and Figure S4)

#### 3) Realistic SE-EPI field maps

We calculated two metrics based on the SE-EPI field maps, including the SE-EPI based mean squared difference between different PE directions (EPI-MSD; lower values indicate better performance), and the SE-EPI based structural similarity (EPI-SS; local cross-correlation between corrected SE-EPI and T2w images with window size of 3; higher values indicate better performance).

#### 4) Multicenter dataset

To evaluate the effectiveness of our method in reducing multicenter effects, we calculated the coefficient of variation (CV) and the intraclass correlation coefficient (ICC). A lower CV indicates less variability for brain measurements of the same subject across centers, while a higher ICC indicates greater consistency. Together, a lower CV and a higher ICC suggest reduced multicenter effects.

We used the one-sided Wilcoxon rank test on these quantitative metrics for statistical comparisons between different methods. The detailed calculations and statistical results of all metrics are described in SI-5 and SI-7, separately.

### 3.3 Baseline methods

The baseline methods were chosen based on three criteria: 1) demonstrating excellent SAC performance in previous studies; 2) being compatible with the specific EPI protocol design; 3) having publicly available code. For inverse-PE datasets, we compared SACNet with the FSL Topup (Andersson et al., 2003) and S-Net (Duong et al., 2020b) approaches. For the single-PE datasets, we compared SACNet with the Fieldmap method included in FSL software and a widely adopted deep-learning image registration baseline, namely, VoxelMorph (Balakrishnan et al., 2019). Notably, we constrained the deformation field along the PE direction in VoxelMorph.

To evaluate the effectiveness of prior neuroanatomical information, we considered three configurations of SACNet by inputting only paired b0 images (SACNet(wos)) or inputting structural images with or without paired b0 images (SACNet(T1w) for T1w image input and SACNet(T2w) for T2w image input). For the simulation data, we only trained SACNet(wos) and SACNet(T2w), as the T1w image is not provided. For the HCP and HCP-D datasets, all configurations were trained. For the dHCP dataset, we only trained SACNet(wos) and SACNet(T2w), as T1w images were not successfully acquired for numerous neonates. For the CBD datasets, we trained SACNet(T1w) and SACNet(T2w) without paired b0 image inputs since these datasets only include single-PE b0 images. For the multicenter dataset, we only trained SACNet(wos) and SACNet(T1w), as this dataset includes no T2w structural images.

### 3.4 Implementations

The proposed method was implemented in Python using the PyTorch software library (Paszke et al., 2019). Our model was trained and tested on a Linux workstation equipped with an Intel Xeon Gold 6258R CPU and a 48 GB GTX Quadro RTX 8000 GPU. We employed the Adam optimizer (Kingma and Ba, 2014) with a learning rate of 1e-4 for optimization. The specific training and inference configurations for each dataset are detailed in SI-3.

## 4. Results

### 4.1 SAC performance on simulated dMRI dataset

We initially evaluate the SACNet using the public simulated dMRI brain scans, as it provide the undistorted dMRI images that can serve as a gold standard. The visual representation of the correction performance and error distribution of different methods are given in Fig. 3. Comparing to Topup and S-Net, SACNet approaches exhibited more recovery details and lower correction errors particularly in the orbitofrontal areas for all 3 types of images including b0, FA and fieldmap. Quantitative analyses (1^st^ row of Table 2) further revealed that SACNet models (both SACNet(wos) and SACNet(T2w)) significantly outperformed Topup and S-Net in all metrics including b0-Err, FA-Err and FM-Err for brain areas with severe distortions (voxels with distortions ≥ 10 mm, for whole brain voxels and other distortion level, see Supplementary Figure S5). SACNet also substantially reduced the unnatural folding compared to Topup and S-Net (NFV value, Topup: 434; S-Net: 136; SACNet: 24-27). All statistical details are shown in Supplementary Table S2.

**Table 2.**
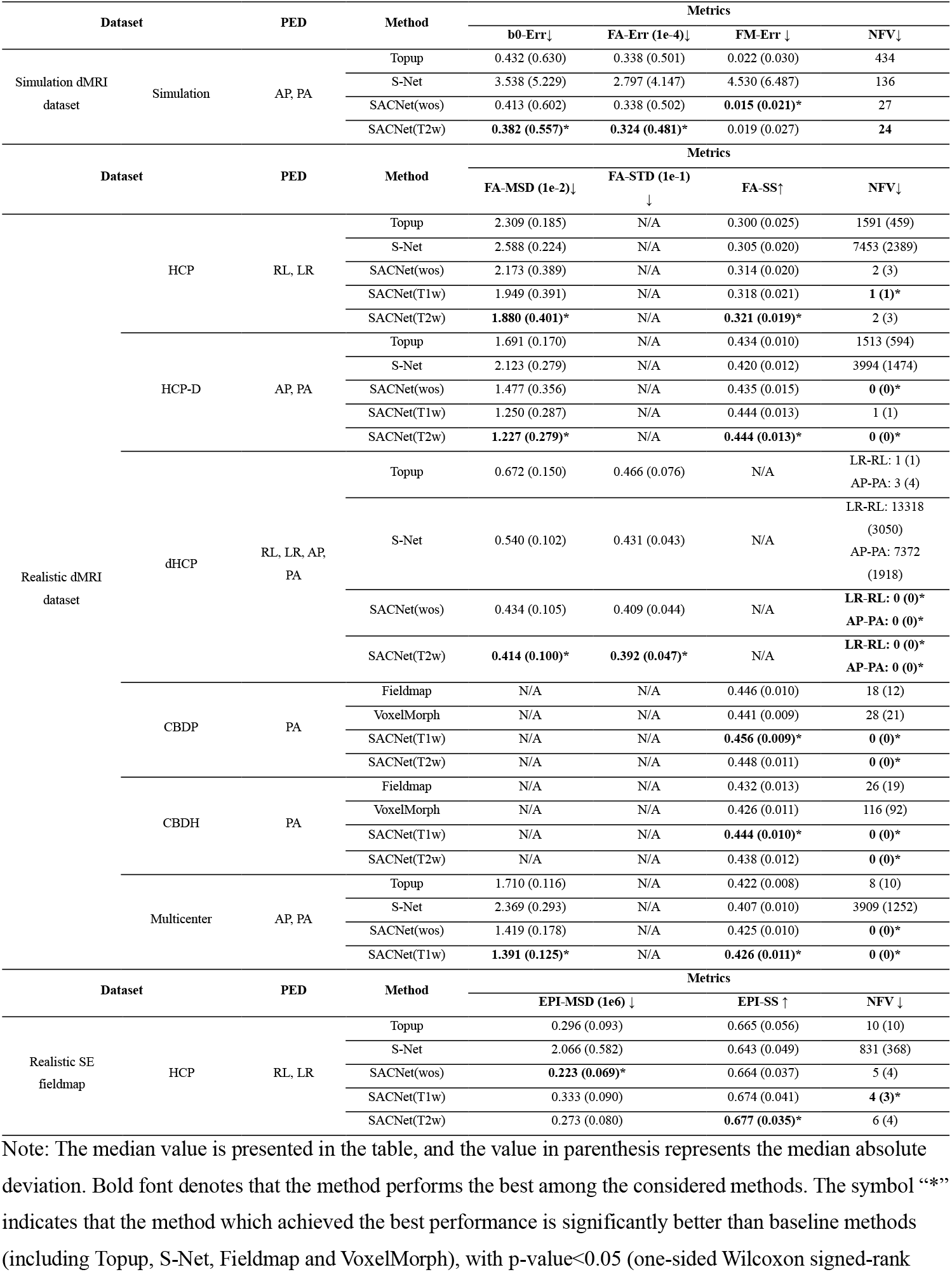
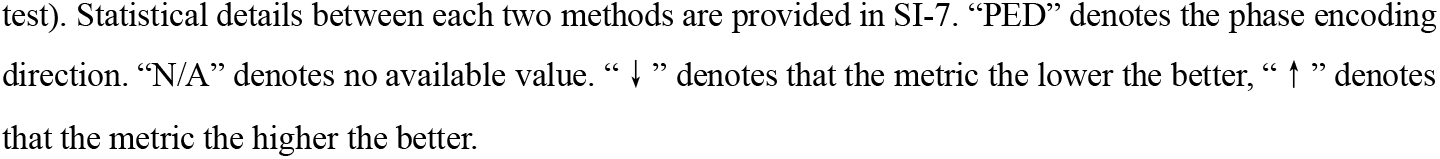
The quantitative performance of SAC results on each dataset.

**Fig. 3.**
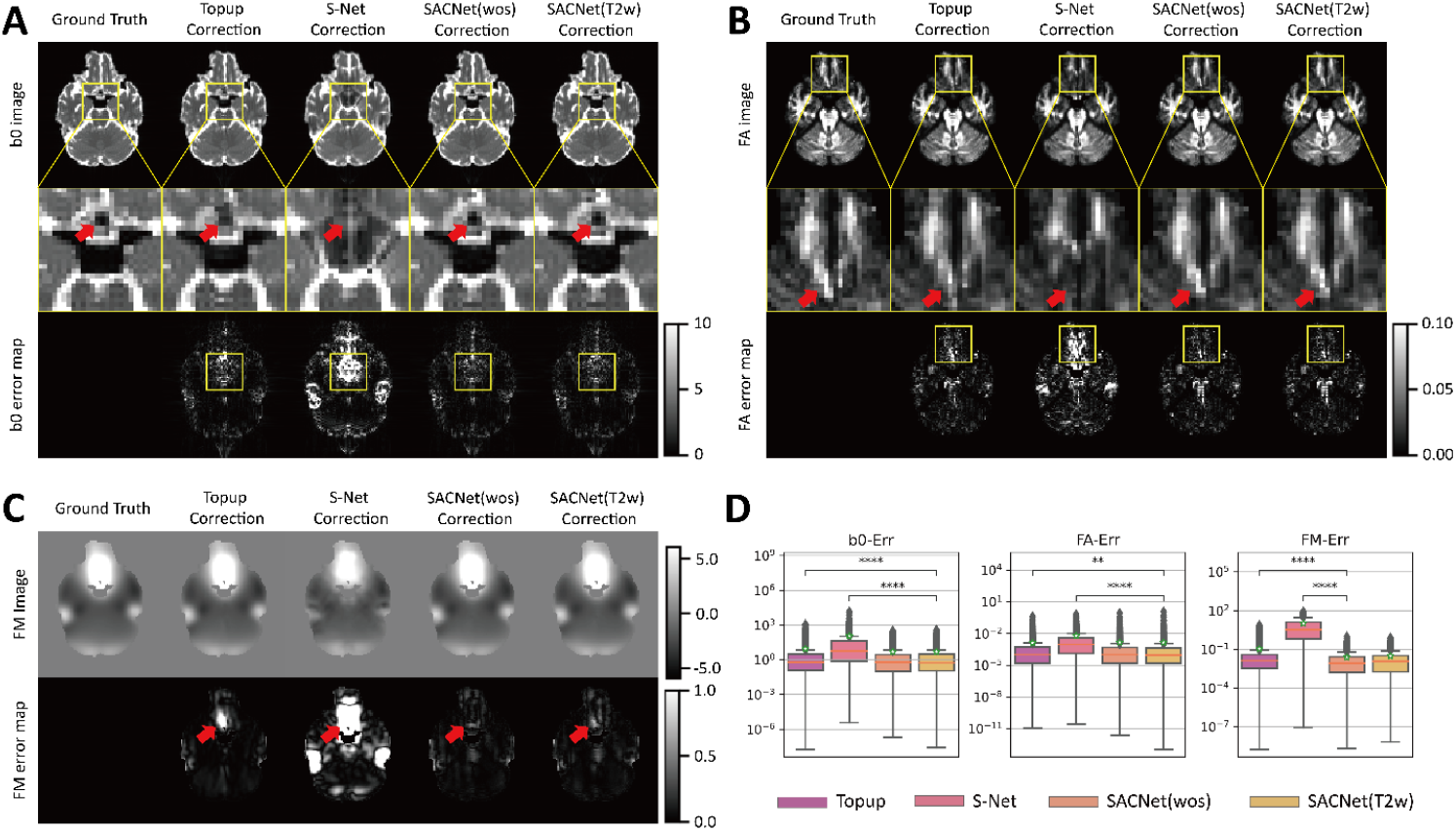
Visualization and quantitative comparation of different SAC methods on simulated dMRI images. The visual view of SAC on b0 images, FA images and field map images across methods are shown in (A), (B) and (C), respectively. The ground truth and corrected images across methods are shown in columns. The raw views, the magnified views and the errors maps are shown in rows. (D) Results of quantitative analyses across methods on brain areas with severe distortions (voxels with distortions ≥10 mm) are depicted in boxplots. FM denotes fieldmap. The green star indicates the mean value, and the coral horizontal line indicates the median value. We marked the significance of group differences between the best-performing model and baseline models. “**” denotes that 0.001 < p ≤ 0.01, and “****” denotes that p ≤ 0.0001.

### 4.2 SAC performance on the inverse-PE adult dataset

For the evaluation on realistic dMRI brain images, we first employed 380 adult brain scans with inverse-PE from HCP dataset (300 scans were used for training, 40 for validation, and 40 for testing). By visualizing the corrected b0 images and FA maps (Fig. 4A) in different approaches, we found that SACNet models bring more construction details, particularly in the frontal cortex, compared to Topup and S-Net. Quantitative comparisons further revealed that the SACNet(T1w) and SACNet(T2w) models significantly outperformed Topup and S-Net in both FA-MSD and FA-SS metrics (all p<0.001, Wilcoxon rank test), achieving improvements of up to 27.4% and 6.5%, respectively (Fig. 4B, 2^nd^ row of Table 2). Of note, the SACNet(T1w) and SACNet(T2w) models achieved significantly lower FA-MSD and higher FA-SS values than the SACNet(wos) model (Fig. 4B, all p<0.001, Wilcoxon rank test), highlighting the importance of incorporating prior neuroanatomical information. Meanwhile, all SACNet variants substantially reduced the NFV metric (Topup: 1591; S-Net: 7453; SACNet: ≤2). Statistical details are shown in Supplementary Table S3.

**Fig. 4.**
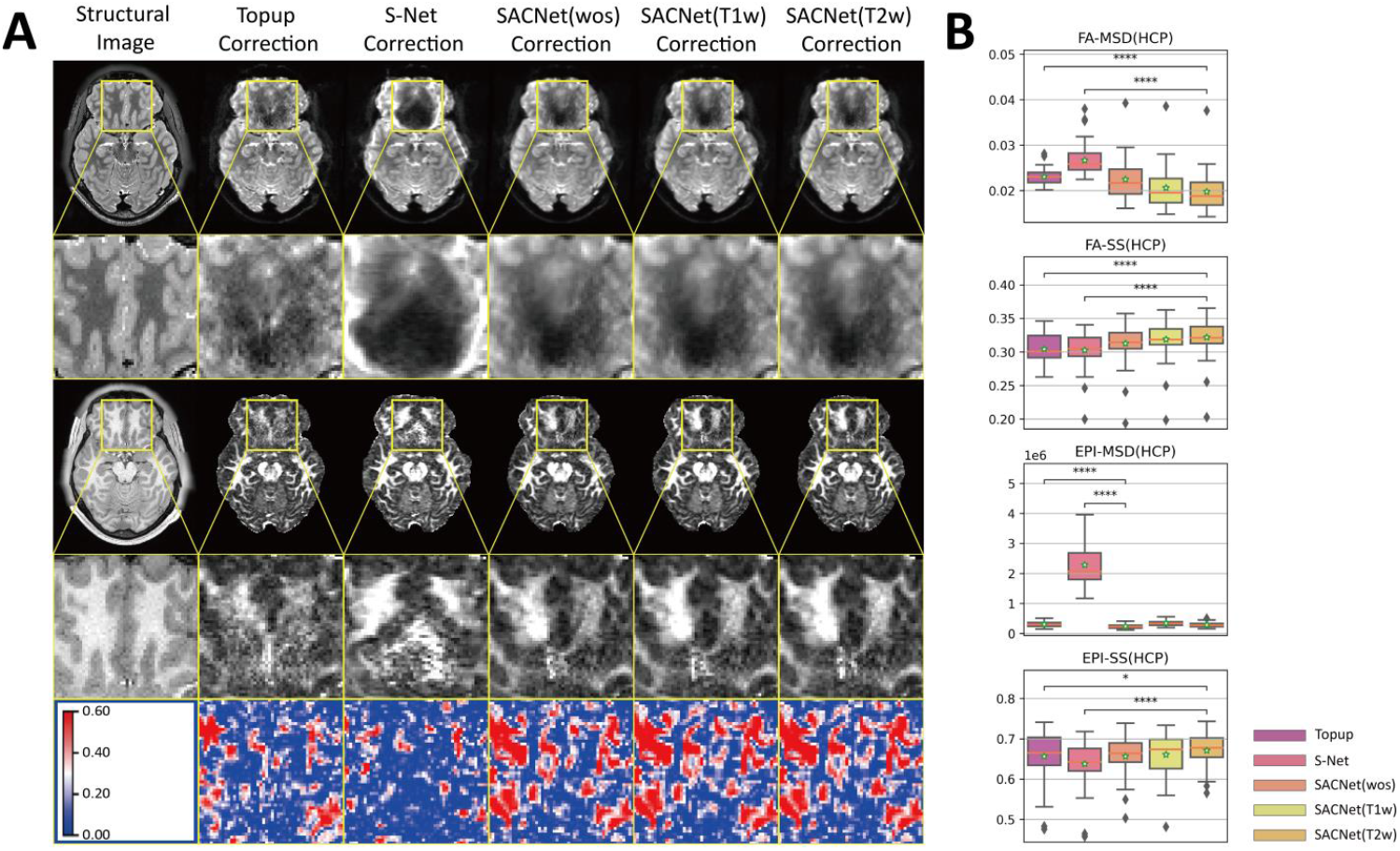
Visualization and quantitative comparison of SAC methods on inverse-PE dMRI images (adult brain). (A) Visualization on SAC results of b0 and FA images, separately. The structural images and corrected dMRI images across different methods are presented in columns. For each method, the raw views and the magnified views are shown, sequentially. The last row presents FA-SS maps (local cross-correlation with a window size of 3 between the FA and T1w images), with warmer colors indicating higher structural similarity. (B) Quantitative results across methods are depicted in boxplots. The green star indicates the mean value, and the coral horizontal line indicates the median value. We marked the significance of group differences between the best-performing model and baseline models. “*” denotes that 0.01 < p ≤ 0.05, and “****” denotes that p ≤ 0.0001.

### 4.3 SAC Performance on the inverse-PE developmental datasets

We further conducted experiments on inverse-PE developmental brain images, using 644 children MR scans from the HCP-D dataset (544 scans were used for training, 50 for validation and 50 for testing) and 444 neonatal brain scans from the dHCP dataset (364 scans were used for training, 40 for validation and 40 for testing). For both the HCP-D and dHCP datasets, our approach demonstrated superior correction quality at frontal cortical boundaries compared to the Topup and S-Net (HCP-D, Fig. 5A; dHCP, Fig. 5B). Quantitative analyses confirmed that SACNet(T2w) significantly outperformed baseline methods in terms of FA-MSD, FA-SS and NFV metrics for HCP-D dataset, and in terms of FA-MSD, FA-STD and NFV metrics for dHCP dataset (Fig. 5C and the 3^rd^ and 4^th^ rows of Table 2, all p<0.001, Wilcoxon rank test). Statistical details were provided in Supplementary Table S4 and S5.

**Fig. 5.**
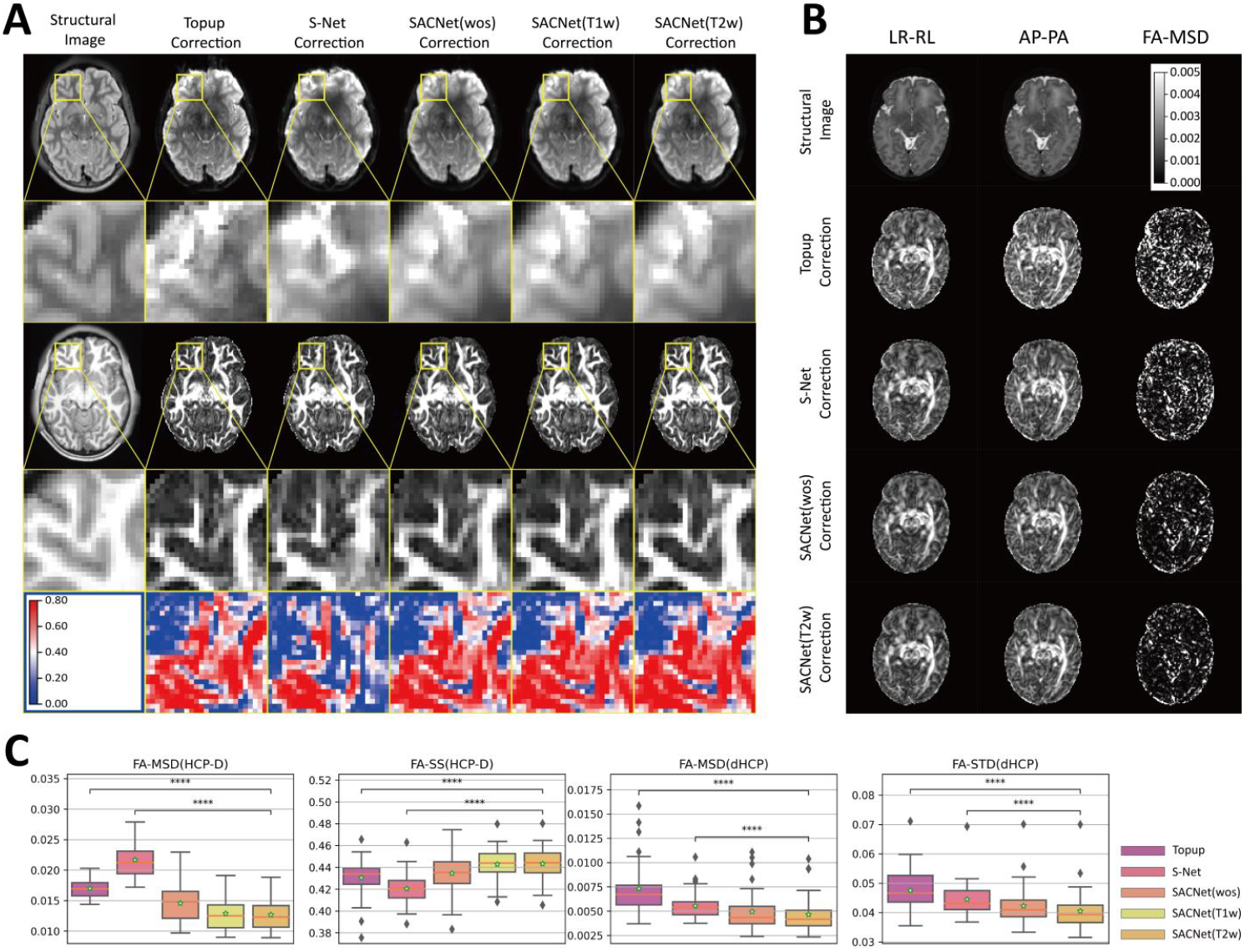
Visualization and quantitative comparison of SAC methods on inverse-PE dMRI images (pediatric brain). (A) Visualization on SAC results of b0 and FA images for HCP-D datasets (adolescents). The structural images and corrected dMRI images across different methods are presented in columns. For each method, the raw views and the magnified views are shown, sequentially. The last row presents FA-SS maps, with warmer colors indicating higher structural similarity. (B) Visualization on SAC results of FA images for dHCP dataset (neonates). The structural images and corrected FA images across different methods are presented in rows. For each method, the views of different PE-encoding approaches are shown, sequentially. The last column presents FA-MSD maps, with brighter colors indicating higher variability across pair-images. (C) Quantitative results across methods are depicted in boxplots for HCP-D and dHCP datasets, respectively. The green star indicates the mean value, and the coral horizontal line indicates the median value. We marked the significance of group differences between the best-performing model and baseline models. “****” denotes that p ≤ 0.0001.

### 4.4 SAC Performance on the single-PE developmental dataset

To evaluate the performance of SACNet on single-PE brain images with dual-echo field maps, we first employed 322 children brain scans from the CBDP dataset (with 242 subjects for training, 40 for validation and 40 for testing). Quantitatively, SACNet(T1w) and SACNet(T2w) both achieved significantly higher FA-SS values and lower NFV metrics compared to FieldMap and VoxelMorph (Fig. 6B and the 5^th^ row Table 2, all p<0.001, Wilcoxon rank test). To further evaluate the generalization of SACNet across centers, we employed 134 children brain scans from another independent subset of CBD project (the CBDH dataset). We directly evaluated the models trained with images of CBDP dataset on the CBDH dataset. We observed high consistent results as that in the CBDP dataset (visual representation in Fig.6A, quantitative results shown in Fig. 6B and the 6^th^ row of Table 2), showing that the SACNet still significantly outperformed FieldMap and VoxelMorph methods (all p<0.05, Wilcoxon rank test). Detailed statistical results are provided in Supplementary Table S6.

**Fig. 6.**
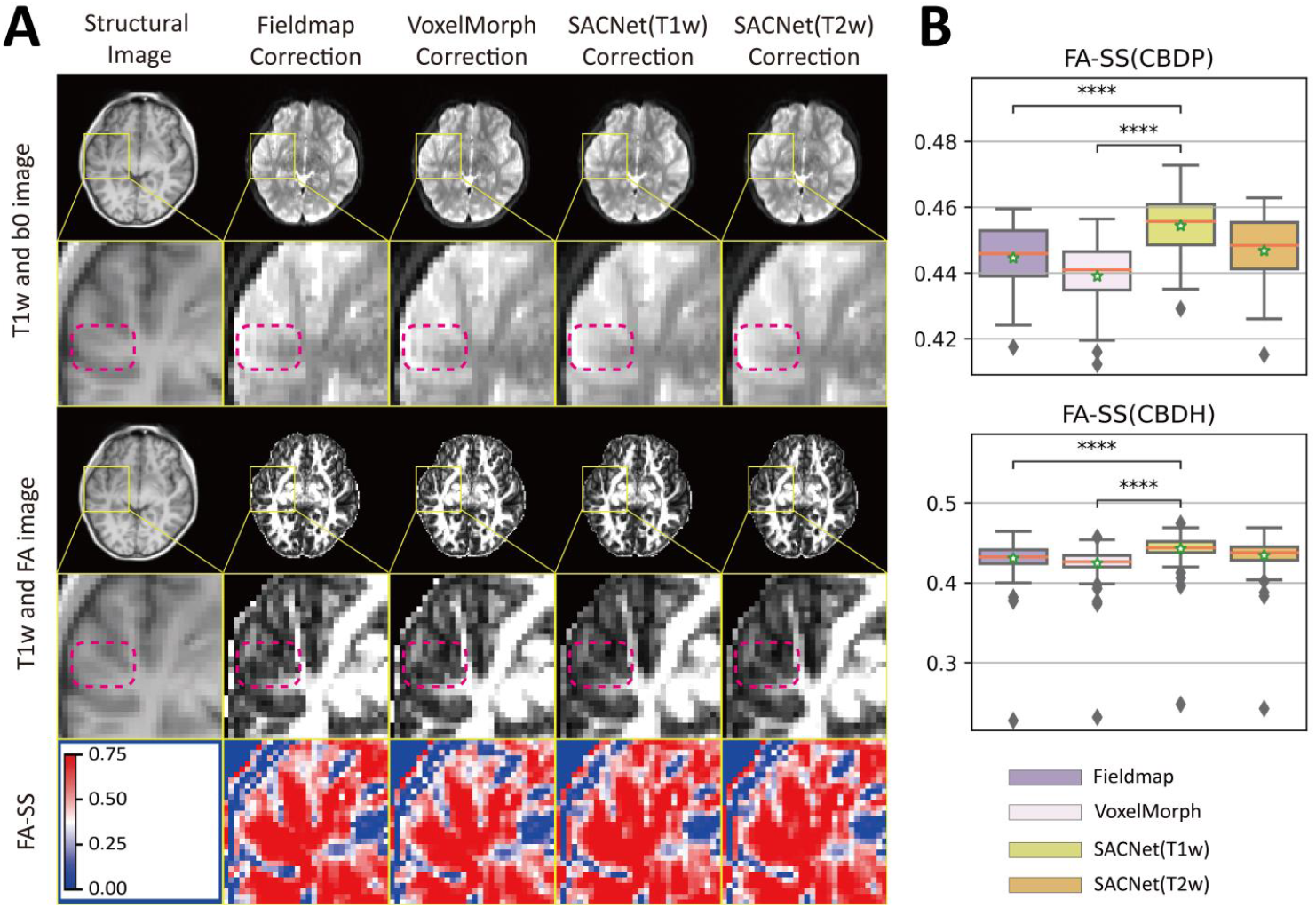
Visualization and quantitative comparison of SAC methods on single-PE dMRI images (developmental brain). (A) Visualization on SAC results of b0 and FA images for CBD datasets. The structural image, uncorrected dMRI images and corrected dMRI images across different methods are presented in columns. For each method, the raw views and the magnified views are shown, sequentially. The last row presents FA-SS maps, with warmer colors indicating higher structural similarity. (B) Quantitative results across methods are depicted in boxplots for CBD datasets. The green star indicates the mean value, and the coral horizontal line indicates the median value. We marked the significance of group differences between the best-performing model and baseline models. “****” denotes that p ≤ 0.0001.

### 4.5 SAC Performance on the SE fieldmaps for fMRI images

Recent neuroimaging projects (such as HCP and BCP project) began to collect an additional SE-EPI scans with inverse PE directions to correct SAs. Such SE-EPI images provide clear anatomical structural details and can be further extended on fMRI brain scans. To evaluate the generalization of our model on such design. We employed SE-EPI field maps from fMRI scans of 379 subjects in HCP dataset. Quantitative results revealed that SACNet(T2w) significantly outperformed both the Topup and S-Net methods in EPI-MSD and EPI-SS metrics (all p<0.05, Wilcoxon rank test) as shown in Fig. 4B and the last row of Table 2. All statistical details are shown in Supplementary Table S3.

### 4.6 SAC Performance on Multicenter dataset with traveling subjects (inverse-PE images)

Finally, we evaluated the performance of the SACNet models with a public multicenter dataset that includes MR images from three traveling subjects collected across 10 sites. To assess whether our model could still achieve excellent SAC performance by only fine-tuning with few images, we employed the models that have been trained with the HCP-D dataset and used three brain scans from one site in the Multicenter dataset to conduct fine-tuning (detailed fine-tuning strategy were provided in SI-6). Quantitative comparisons show that the SACNet(T1w) model significantly outperformed the Topup and S-Net in terms of FA-MSD and FA-SS (Fig. 7A and 7^th^ row of Table 2, all p<0.005, Wilcoxon rank test). Detailed statistical results are provided in Supplementary Table S7.

**Fig. 7.**
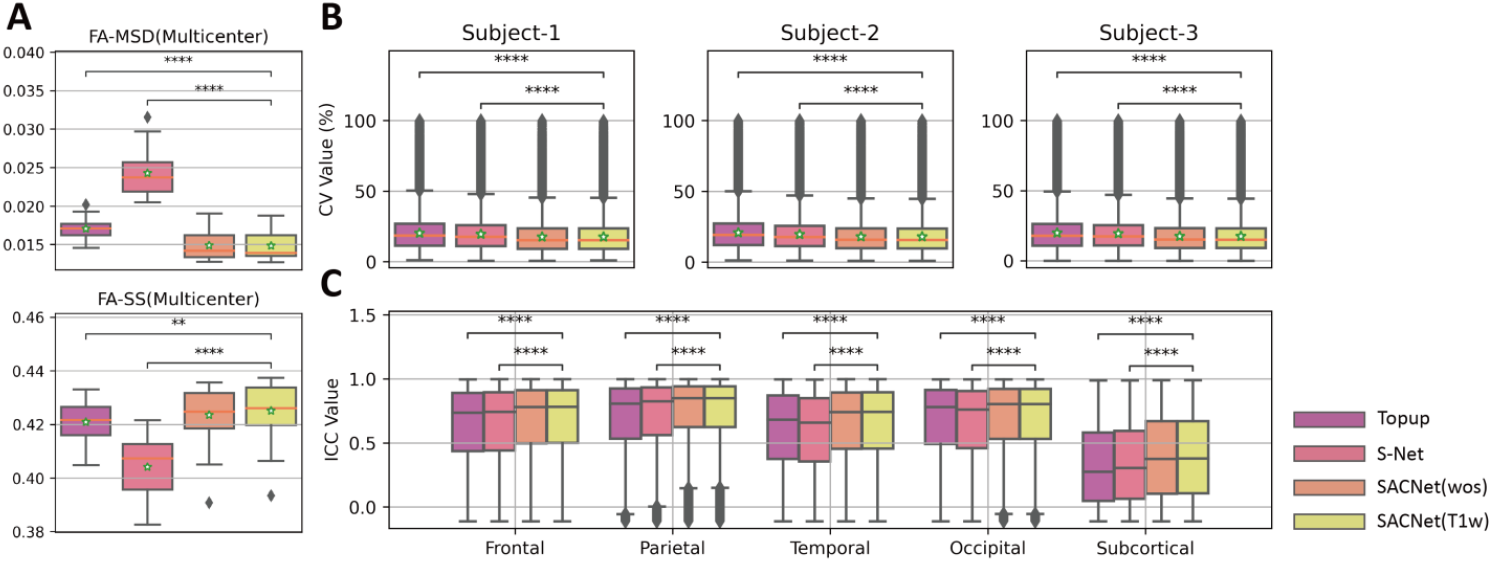
Evaluations on multi-center dMRI images. (A) The SAC performance of each method. (B) Boxplots of the CV distribution for each method in three traveling subjects. (C) Boxplots of the ICC distribution in each brain lobe for each method. The green star indicates the mean value, and the coral horizontal line indicates the median value. We marked the significance of group differences between the best-performing model and baseline models. “**” denotes that 0.001 < p ≤ 0.01, and “****” denotes that p ≤ 0.0001.

To further assess whether our SAC model could reduce multicenter effects. We calculated the CV and ICC metrics with lower CVs or higher ICCs indicating smaller multicenter noise residuals. The SACNet(T1w) achieved significantly lower CV and higher ICCs than Topup and S-Net (Fig. 7B and 7C, all p<0.001, Wilcoxon rank test). Notably, SACNet(T1w) obtained significantly higher ICC values than SACNet(wos) in the frontal and temporal lobe that were severely affected by SAs (Fig. 7C, all p<0.001, Wilcoxon rank test). This suggests that incorporating prior neuroanatomical information in SAs correction is valuable for reducing the multicenter effects in severely distorted areas. Detailed statistical results are provided in Supplementary Tables S8 and S9.

### 4.7 Ablation tests

We separately assessed the effectiveness of the differentiable EPI warp (DEW) module, diffeomorphism preservation function (DPF), and coarse-to-fine (CTF) training and inference protocols (Section 4.7.1) through ablation studies. Additionally, we evaluated each component loss in the prior neuroanatomical information loss (Section 4.7.2). These experiments were conducted using the HCP dataset, with T2w images serving as the prior neuroanatomical information.

#### 4.7.1 Ablation studies on the DEW module, DPF and CTF protocols

The effects of different combinations of the three components are presented in Table 3. Without the DEW module, the proposed method exhibited the worst SAC performance, characterized by the highest FA-MSD and lowest FA-SS values. Omitting the diffeomorphism preservation function (DPF) led to an increase in the number of folding voxels and a decline in SAC performance decreased. Excluding the coarse-to-fine training and inference (CTF) protocols also resulted in decreased SAC performance, with increased FA-MSD and decreased FA-SS values. Additionally, SACNet required approximately 24 hours for training without the CTF protocols, but only around 12 hours with them.

**Table 3.** The quantitative results of the ablation study based on the use of the DEW module, DPF and CTF protocols.

| DEW | DPF | CTF | Metrics |  |  |
| --- | --- | --- | --- | --- | --- |
|  |  |  | FA-MSD (1e-2) | FA-SS | NFV |
| ✗ | ✓ | ✓ | 2.461 (0.222) | 0.299 (0.016) | 51 (62) |
| ✓ | ✗ | ✓ | 1.901 (0.404) | 0.320 (0.019) | 513 (511) |
| ✓ | ✓ | ✗ | 2.002 (0.349) | 0.321 (0.019) | 6 (8) |
| ✓ | ✓ | ✓ | **1.880 (0.401)** | **0.321 (0.019)** | **2 (3)** |

Note: The median value is presented in the table, and the value in parenthesis represents the median absolute deviation. Bold font denotes that the method performs the best among the considered methods.

These results highlight the importance of each component in the proposed method. We further demonstrated the impact of DPF on image correction in Fig. 8. It is evident that the white matter areas of the corrected b0 image aligns more closely with corresponding T2w image (Fig. 8A) after applying DPF (Fig. 8C) compared to before applying DPF (Fig. 8B1 and 8B2).

**Fig. 8.**
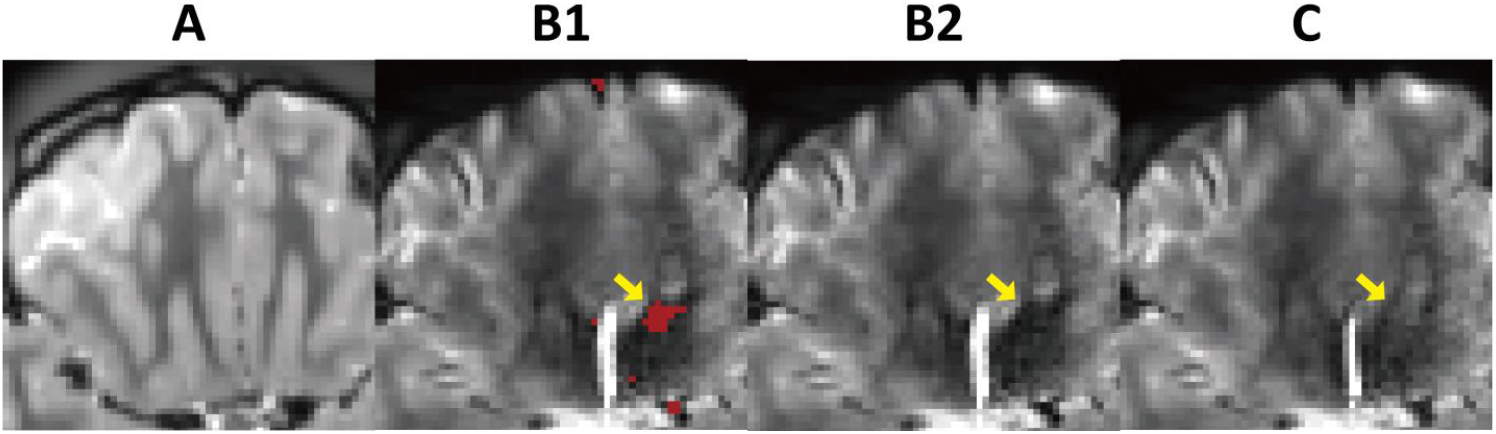
The impact of DPF on image correction. (A) showcases the T2w image. (B1) and (B2) present the corrected b0 images obtained using SACNet(T2w) without DPF, with B1 highlighting folding voxels in red for clarity. (C) presents the corrected b0 image obtained by SACNet(T2w) with DPF.

#### 4.7.2 Ablation studies on each component loss in ℒ_*struct*_

ℒ_*struct*_contains the overall shape structural similarity loss ℒ_*str*−*overall*_ and the pairwise structural similarity loss ℒ_*str*−*pair*_. Table 4 shows that the SAC performance of the model depended on each component loss in ℒ_*struct*_. The best performance was achieved by leveraging both ℒ_*str*−*overall*_ and ℒ_*str*−*pair*_ in the network optimization. Furthermore, ℒ_*str*−*overall*_ and ℒ_*str*−*pair*_ both improved the SAC performance independently.

**Table 4.** The quantitative results of the ablation study of **ℒ**_***str***_.

| $\mathcal{L}_{str-overall}$ | $\mathcal{L}_{str-pair}$ | Metrics | | |
| --- | --- | --- | --- | --- |
|  |  | FA-MSD (1e-2) | FA-SS | NFV |
| ✗ | ✗ | 2.173 (0.389) | 0.314 (0.020) | **2 (3)** |
| ✗ | ✓ | 1.984 (0.387) | 0.318 (0.020) | **2 (3)** |
| ✓ | ✗ | 1.937 (0.384) | 0.319 (0.020) | 5 (7) |
| ✓ | ✓ | **1.880 (0.401)** | **0.321 (0.019)** | **2 (3)** |

Note: The median value is presented in the table, and the value in parenthesis represents the median absolute deviation. Bold font denotes that the method performs the best among the considered methods.

### 4.8 Runtime analysis

Table 5 shows the running time for estimating the inhomogeneity field of single subject based on each dataset for three SA correction methods: SACNet and two conventional methods, Fieldmap and Topup. The results show that SACNet is significantly more efficient than the conventional methods. These results indicate that SACNet has a significant advantage in processing large-scale datasets due to its ultrafast computational speed compared to those of conventional methods.

**Table 5.** Running time (in seconds).

| Method | HCP | HCP-D | dHCP | CBD | Multicenter |
| --- | --- | --- | --- | --- | --- |
| Fieldmap | N/A | N/A | N/A | ~4 hours | N/A |
| Topup | 3061 | 1740 | 986 | N/A | 1320 |
| SACNet (CPU) | 3.87 | 3.57 | 1.30 | 0.89 | 3.22 |
| SACNet (GPU) | 2.17 | 1.52 | 0.73 | 0.46 | 1.30 |

Note: N/A denotes no available value. Here we used the SACNet(T2w) model to evaluate the inference running time.

## 5. Discussion and conclusion

We proposed an unsupervised multiscale convolutional registration network (SACNet) to remove SAs in brain EPI images. This model could generate diffeomorphic inhomogeneity fields based on either inverse-PE or single-PE images and employ prior neuroanatomical constraints from additional T1w or T2w images. Extensive experiments on both simulated and realistic datasets showed that our SACNet not only outperformed most popular conventional correction methods, such as Topup and Fieldmap, but also surpassed deep-learning based methods, such as S-Net and VoxelMorph. Furthermore, by a fine-tuning strategy with few samples in a multicenter dataset with traveling subjects, our model showed better SAC performance and lower multicenter effects than the Topup and S-Net approaches. Our model reduced the time cost of distortion correction from tens of minutes (conventional iterative approaches) to a few seconds while maintaining state-of-the-art SAC performance and robustness. We hope such model could largely facilitate the integration of multicenter MR datasets in future neuroimaging studies.

### 5.1 Applications to large-scale neuroimaging studies

Recent brain neuroimaging investigations have entered the era of “big data” (Bethlehem et al., 2022; Landhuis, 2017; Rutherford et al., 2022; Sejnowski et al., 2014; Xia and He, 2017) by integrating tens of thousands of image scans acquired at multiple centers. We emphasize that our SACNet approach is well suited for large-scale neuroimaging studies involving many individual scans for several reasons. First, we provided a range of models that have been pretrained based on diverse datasets with different ages and acquisition protocols, enabling users to fine-tune the models with only a few images according to their needs and to achieve excellent SAC in their own datasets (Section 4.1-4.6). Although recent cohort projects have used uniform EPI phase encoding protocols, many legacy datasets were acquired with various EPI protocols or even no EPI artifact correction sequences. Considering the high cost of acquiring human brain MR images, the utilization of existing databases is highly valuable. Second, our SACNet model effectively reduced the potential multicenter effects related to SAs (Section 4.6). Recent approaches for multicenter effect correction in brain MR images have received much methodological attention. Our model significantly reduced multicenter noise without using additional correction algorithms, highlighting the necessity of considering SAC in current multicenter correction frameworks (Tian et al., 2022). Third, compared with conventional iterative optimization methods, SACNet can process many images from multiple subjects at fast speeds due to its ultrafast inference time (Section 4.8). Notably, SACNet does not require a large amount of CPU memory (approximately 3000 MB), making it convenient for batch processing in computing clusters. Finally, compared with existing deep-learning based SAC studies, for which only the source code is available, we improved the usability of our model by developing a comprehensive interface of existing dMRI processing pipelines (Glasser et al., 2013) and utilizing the containerization technique for easy installation.

### 5.2 Effective network designs and integrated loss constraints in SACNet enable excellent SAC performance

A common deep-learning based brain registration framework is insufficient for solving the SAC problem. Thus, we proposed several key designs to ensure the high quality of the generated inhomogeneity field and carefully evaluated their effectiveness. First, introducing the DEW module effectively eliminates SAs by correcting geometric and intensity distortions, thus promoting network convergence, as shown in the comparison between the first and last rows in Table 3. Second, the diffeomorphism preservation function (DPF) constrains the inhomogeneity field within the diffeomorphic space, reducing invalid voxels (negative intensity values and folding patterns) during differentiable EPI warp calculation, thus preventing overfitting, as seen in the second and last rows in Table 3. Fig. 8 visually highlights the improved SAs correction with DPF. Instead of using the velocity integration method to generate diffeomorphic deformations, which requires multiple integration steps and longer training time, we employed DPF to directly generate diffeomorphic inhomogeneity field. Third, due to the severe SAs at temporal and frontal cortical boundaries (especially at the temporal pole and orbitofrontal cortex), anatomical morphologies within certain brain locations were barely conserved. Thus, it is imperative to incorporate prior neuroanatomical information to obtain excellent correction results. Previous methods that used structural images as additional inputs to the network were not sufficient to obtain good morphological images since these methods only provide information features and do not contribute to the loss function calculation (Hu et al., 2020; Schilling et al., 2020). To address this issue, we carefully designed a prior neuroanatomical information loss function, ℒ_*struct*_, that was optimized for SAC by incorporating gradient-based information from structural images. This function ℒ_*struct*_includes two components: ℒ_*str*−*pair*_ and ℒ_*str*−*overall*_. ℒ_*str*−*pair*_ was used to align the corrected image pair 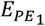 and 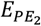 to the structural image *I*_*struct*_in a pairwise manner (third row vs. last row in Table 4), and ℒ_*str*−*overall*_ improved the overall structural alignment between the final corrected image *E*_*final*_ and *I*_*struct*_(second row vs. last row in Table 4). Moreover, the choice of an intensity-irrelevant structural metric, the NGF, allows users to use either T1w or T2w images as input neuroanatomical information, thereby improving the compatibility of SACNet for different types of clinical datasets. Last, the well-designed CTF training and inference protocols adopted in SACNet accelerated the training process and improved model convergence (third row vs. last row in Table 3), and similar strategies have been broadly deployed in conventional SAC methods (Bhushan et al., 2015; Duong et al., 2020a; Irfanoglu et al., 2015; Ruthotto et al., 2012).

### 5.3 Comparison with deep-learning based methods

Previous studies have proposed several deep-learning based registration approaches to address the SAC problem. For example, Bian et al. proposed correcting distortions by registering distorted b0 images in a single-PE direction to T1w images through the VoxelMorph backbone by optimizing the mutual information (MI) loss (Bian et al., 2023). However, this approach is limited to single-PE type data and was not compared with other methods designed specifically for the SAC problem. Duong et al. and Zahneisen et al. predicted inhomogeneity fields to remove SAs with 3D and 2D CNNs, respectively (Duong et al., 2020b; Zahneisen et al., 2020). However, the performance of these models is limited, as the models either ignore the intensity distortion problem or treat each volume slice as an independent example for training, resulting in inadequate SAs correction or inconsistent alignment between slices. Other methodological approaches for solving the SAC problem have also been developed. Several studies have used image generation approaches for SAC tasks. For example, Hu et al. and Ye et al. used high-resolution distortion-free point spread function encoded EPI (PSF-EPI) data as undistorted ground truth data for CNN training (Hu et al., 2020; Ye et al., 2023). Schilling et al. synthesized undistorted b0 images with U-Net or generative adversarial networks (GANs) and then entered both the “synthesized” and “real” b0 images as input into Topup to remove SAs (Schilling et al., 2020; Schilling et al., 2019). These supervised approaches need ground truth images as learning targets, which largely depend on the feature distribution of the training images. This may lead to difficulty when facing brain images with heterogeneous appearance, such as those of neonatal brain scans. Moreover, when these approaches are applied to a new neuroimaging dataset, acquiring undistorted images for fine-tuning can often be a costly endeavor. In contrast to image generation-based methods, unsupervised registration-based methods, such as our approach, require no ground truth labels. Interestingly, Qiao et al. proposed the distortion correction network (DrCNet) by feeding fiber orientation distribution (FOD) information into U-Net and successfully corrected residual distortions that could not be eliminated by Topup (Qiao and Shi, 2021). Compared to DrCNet, our SACNet method can be applied not only to dMRI data but also to fMRI data by using additionally collected spin-echo field maps (Section 4.5). Moreover, the performance of SACNet may be further enhanced by incorporating rich diffusion-based information, such as DWIs and FODs, into the integrated loss function.

### 5.4 Limitations and future directions

Several issues in this study should be considered. First, although the additional neuroanatomy priors improve the SAC performance of our SACNet, it may be worthwhile to explore the further employment of white matter information from dMRI data itself (Irfanoglu et al., 2015; Qiao and Shi, 2021; Qiao et al., 2019). Second, the presence of other nonnegligible artifacts, such as eddy current-induced distortions and intrasubject movements, in dMRI data should be acknowledged (Andersson and Sotiropoulos, 2016). It would be interesting to develop a deep-learning based tool in conjunction with SACNet to address these artifacts. Third, registration-based methods may become unsatisfactory in 7T MR images due to severe signal loss issues. Given that the deep generative model (DGM) has shown the ability to capture complex distributions of real 7T data (Nie et al., 2018), combining the DGM with SACNet could be a promising approach. We hope that SACNet can offer a general framework for SAC task in multicenter datasets with top-ranking performance, robust output and efficient computational speed, which could facilitate a wide variety of future brain studies using large-scale multicenter neuroimaging datasets.

## Data availability

The datasets from the Human Connectome Project (HCP) and the Lifespan Human Connectome Project Development (HCP-D) are available at https://www.humanconnectome.org. The dataset from the Developing Human Connectome Project (dHCP) is available at https://www.developingconnectome.org. The Multicenter dataset is available at https://doi.org/10.6084/m9.figshare.8851955.v6. The simulation dataset is available at https://www.nitrc.org/projects/diffusionsim. Raw imaging data from the Children School Functions and Brain Development Project (CBD) is available from the corresponding authors upon reasonable request.

## Code availability

The source code that implements our software is available at https://github.com/RicardoZiTseng/SACNet.

## Acknowledgments

The study was supported by the National Natural Science Foundation of China (Nos. 31830034, 82021004, 81801783, 82102139 and 82202243) and the Changjiang Scholar Professorship Award (T2015027). We are grateful to the Developing Human Connectome Project (DHCP), the Human Connectome Project (HCP) and the Lifespan Human Connectome Project Development (HCPD). HCP imaging data were provided by the Human Connectome Project, WU-Minn Consortium (Principal Investigators: David Van Essen and Kamil Ugurbil; 1U54MH091657) funded by the 16 NIH Institutes and Centers that support the NIH Blueprint for Neuroscience Research and by the Mc-Donnell Center for Systems Neuroscience at Washington University.

The Lifespan HCP imaging data were supported by the National Institute of Mental Health of the National Institutes of Health under Award Number U01MH109589 and by funds provided by the McDonnell Center for Systems Neuroscience at Washington University in St. Louis. The content is solely the responsibility of the authors and does not necessarily represent the official views of the National Institutes of Health.

## Author contributions

Z.L.Z., T.D.Z., and Y.H. designed the research; W.W.M., Y.P.W., R.C., H.B.Z., S.P.T., J.H.G., S.Z.Q., Q.Q.T., H.J.H., S.T., Q.D., and Y.H. collected the imaging dataset of CBD project; J.Y.Z., X.Y.L., L.L.S., Y.H.Z., and Y.H. provided the methodological instruction; Z.L.Z., and T.D.Z. performed the data analysis; Z.L.Z., and T.D.Z. wrote the paper; Z.L.Z., T.D.Z., and Y.H. revised the paper.

