## Supplemental Information Appendix for "SACNet: A Multiscale Diffeomorphic Convolutional Registration Network with Prior Neuroanatomical Constraints for Flexible Susceptibility Artifact Correction in Echo Planar Imaging"

*** Corresponding author:**

#### SI-1. Jacobian determinant of the transformation $\boldsymbol{p} \rightarrow \boldsymbol{p} + \boldsymbol{B}(\boldsymbol{p})\boldsymbol{v}$

In this section, we prove the Jacobian determinant of the transformation $\boldsymbol{p} \rightarrow \boldsymbol{p} + B(\boldsymbol{p})\boldsymbol{v}$. For $\boldsymbol{v} = (1,0,0)$, let $\partial_v B(p)$ denote the directional derivative of field $B$ at point $\boldsymbol{p}$ along the PE direction $\boldsymbol{v}$. The Jacobian determinant of the transformation $\boldsymbol{T}(\boldsymbol{p}): \boldsymbol{p} \rightarrow \boldsymbol{p} + B(\boldsymbol{p})\boldsymbol{v}$ at point $p$ can be calculated as follows:

$$
\begin{aligned}
Jaco_{\boldsymbol{T}(\boldsymbol{p})} &= \frac{\partial(\boldsymbol{p} + B(\boldsymbol{p})\boldsymbol{v})}{\partial \boldsymbol{p}} \\
&= \begin{bmatrix}
\frac{\partial(\boldsymbol{p} + B(\boldsymbol{p})\boldsymbol{v})_x}{\partial p_x} & \frac{\partial(\boldsymbol{p} + B(\boldsymbol{p})\boldsymbol{v})_x}{\partial p_y} & \frac{\partial(\boldsymbol{p} + B(\boldsymbol{p})\boldsymbol{v})_x}{\partial p_z} \\
\frac{\partial(\boldsymbol{p} + B(\boldsymbol{p})\boldsymbol{v})_y}{\partial p_x} & \frac{\partial(\boldsymbol{p} + B(\boldsymbol{p})\boldsymbol{v})_y}{\partial p_y} & \frac{\partial(\boldsymbol{p} + B(\boldsymbol{p})\boldsymbol{v})_y}{\partial p_z} \\
\frac{\partial(\boldsymbol{p} + B(\boldsymbol{p})\boldsymbol{v})_z}{\partial p_x} & \frac{\partial(\boldsymbol{p} + B(\boldsymbol{p})\boldsymbol{v})_z}{\partial p_y} & \frac{\partial(\boldsymbol{p} + B(\boldsymbol{p})\boldsymbol{v})_z}{\partial p_z}
\end{bmatrix} \\
&= \begin{bmatrix}
1 + \partial_v(B(\boldsymbol{p})) & 0 & 0 \\
0 & 1 & 0 \\
0 & 0 & 1
\end{bmatrix} = 1 + \partial_v(B(\boldsymbol{p}))
\end{aligned}
\qquad (S1)
$$

#### SI-2. Proof of the existence of diffeomorphic solution

In this section, we prove the existence of at least one minimizer of Eq. (5) and that the positive intensity modulations and diffeomorphic space transformation can be ensured by the DPF. This proof uses the framework of direct methods in variational calculus. For more explanation of the underlying theory, please refer to (Evans, 1998). Similar proof can be found in the work of Ruthotto et al. (Ruthotto et al., 2012).

**Theorem 1.** For images $I_{PE_1}, I_{PE_2}, I_{struct} \in C^1(\Omega)$ and regularization parameters $\alpha, \beta, \delta, \gamma_1, \gamma_2 > 0$, assume that the loss $\mathcal{L}(I_{PE_1}, I_{PE_2}, I_{struct}, B)$ expressed by Eq. (5) is finite and that $\mathcal{L}(I_{PE_1}, I_{PE_2}, I_{struct}, \boldsymbol{0}) < \infty$. Then, at least one minimizer $B^*$ of $\mathcal{L}(I_{PE_1}, I_{PE_2}, I_{struct}, B)$ exists in the solution set $\mathcal{S}$:

$$\mathcal{S} = \left\{ B \in W^{1,2}(\Omega) \middle| \frac{1}{|\Omega|} \int_\Omega |B(\boldsymbol{p})| d\boldsymbol{p} \leq K, |\partial_v B(\boldsymbol{p})| < 1 \right\} \qquad (S2)$$

where the mean magnitude of all feasible inhomogeneity field sets is bounded by $K \in [0, +\infty]$, and $W^{1,2}(\Omega)$ denotes the Sobolev space.

**Proof**. We first proved that the inhomogeneity field $B$ belongs to $W^{1,2}(\Omega)$; then, we proved that $\mathcal{L}(I_{PE_1}, I_{PE_2}, I_{struct}, B)$ is coercive and has lower semicontinuity, which ensures the existence of a minimizer $B^*$ in $W^{1,2}(\Omega)$.

**1. Prove that $\boldsymbol{B} \in \boldsymbol{W}^{1,2}(\boldsymbol{\Omega})$.**

$B(\boldsymbol{p})\colon \Omega \to \mathbb{R}$ is a local integrable function, and for any multiple indices $|\alpha| \leq 1$, we have $D^\alpha B \coloneqq \{B, \partial_v B\}$. In addition, $\|B\|_{L^2(\Omega)} < \infty$, $\|\partial_v B\|_{L^2(\Omega)} < \infty$, and thus $D^\alpha B \in L^2(\Omega)$; thus, we have $B \in W^{1,2}(\Omega)$.

**2. Prove the coercivity of $\boldsymbol{\mathcal{L}}\left(\boldsymbol{I}_{\boldsymbol{PE_1}}, \boldsymbol{I}_{\boldsymbol{PE_2}}, \boldsymbol{I}_{\boldsymbol{struct}}, \boldsymbol{B}\right)$.**

According to Eq. (5), we can estimate the lower bound of $\mathcal{L}\left(I_{PE_1}, I_{PE_2}, I_{struct}, B\right)$ as:

$$\mathcal{L}\left(I_{PE_1}, I_{PE_2}, I_{struct}, B\right) \geq \delta \cdot \mathcal{L}_{diff}(B) = \delta \cdot \|\nabla B\|^2_{L^2(\Omega)} \qquad (S3)$$

In addition, according to the Poincaré inequality, we have

$$\|B - B_\Omega\|^2_{L^2(\Omega)} \leq C\|\nabla B\|^2_{L^2(\Omega)}$$

$$\Rightarrow \int_\Omega |B(\boldsymbol{p}) - B_\Omega|^2 d\boldsymbol{p} \leq C \int_\Omega |\nabla \mathrm{B}(\mathbf{p})|^2 d\boldsymbol{p}$$

$$\Rightarrow C\left(\int_\Omega |B(\boldsymbol{p})^2|\, d\boldsymbol{p} - \int_\Omega |B_\Omega|^2\, d\boldsymbol{p}\right) \leq C \int_\Omega |B(\boldsymbol{p}) - B_\Omega|^2 d\boldsymbol{p} \leq \int_\Omega |\nabla B(\mathbf{p})|^2 d\boldsymbol{p}$$

$$\Rightarrow C\left(\|B\|^2_{L^2(\Omega)} - |\Omega|\left(\frac{1}{|\Omega|}\int_\Omega B(\boldsymbol{p})\, d\boldsymbol{p}\right)^2\right) \leq \|\nabla B\|^2_{L^2(\Omega)}$$

$$\Rightarrow C \cdot \|B\|^2_{L^2(\Omega)} + C \cdot \|\nabla B\|^2_{L^2(\Omega)} - C \cdot |\Omega|\left(\frac{1}{|\Omega|}\int_\Omega B(\boldsymbol{p})\, d\boldsymbol{p}\right)^2 \leq (C+1) \cdot \|\nabla B\|^2_{L^2(\Omega)}$$

$$\Rightarrow C \cdot \|B\|^2_{W^{1,2}(\Omega)} - C \cdot |\Omega| K^2 \leq C \cdot \|B\|^2_{W^{1,2}(\Omega)} - C \cdot |\Omega|\left(\frac{1}{|\Omega|}\int_\Omega B(\boldsymbol{p})\, d\boldsymbol{p}\right)^2 \leq \|\nabla B\|^2_{L^2(\Omega)} \quad (S4)$$

Thus, we have:

$$\mathcal{L}\left(I_{PE_1}, I_{PE_2}, I_{struct}, B\right) \geq C \cdot \|B\|^2_{W^{1,2}(\Omega)} - C \cdot |\Omega| K^2 \qquad (S5)$$

**3. Prove the lower semicontinuity of $\boldsymbol{\mathcal{L}}\left(\boldsymbol{I}_{\boldsymbol{PE_1}}, \boldsymbol{I}_{\boldsymbol{PE_2}}, \boldsymbol{I}_{\boldsymbol{struct}}, \boldsymbol{B}\right)$.**

Due to the convexity of the integrand with respect to $\nabla B$ between -1 and 1, the lower semicontinuity of $\mathcal{L}\left(I_{PE_1}, I_{PE_2}, I_{struct}, B\right)$ is ensured.

**4. Prove the existence of a minimizer $\boldsymbol{B}^*$ in $\boldsymbol{W}^{1,2}(\boldsymbol{\Omega})$.**

The coercivity and lower semicontinuity of $\mathcal{L}\left(I_{PE_1}, I_{PE_2}, I_{struct}, B\right)$ yield the existence of a minimizer $B^*$ in $W^{1,2}(\Omega)$.

**5. Prove that the minimizer $\boldsymbol{B}^*$ is diffeomorphic.**

Consider a set with a sufficiently small $\epsilon > 0$, $\mathcal{A}_\epsilon \coloneqq \{\boldsymbol{x} \in \Omega | |\partial_v B^*(\boldsymbol{x})| \geq 1 - \epsilon\}$; then, we have:

$$\mathcal{L}(I_{PE_1}, I_{PE_2}, I_{struct}, \mathbf{0}) \geq \mathcal{L}(I_{PE_1}, I_{PE_2}, I_{struct}, B^*) \geq \delta \int_{S_\epsilon} \phi(1-\epsilon)dx = \delta\phi(1-\epsilon)|\mathcal{A}_\epsilon| \quad (S6)$$

Thus, we have:

$$|\mathcal{A}_\epsilon| \leq \frac{\mathcal{L}(I_{PE_1}, I_{PE_2}, I_{struct}, \mathbf{0})}{\delta\phi(1-\epsilon)} \quad (S7)$$

when $\delta$ is sufficiently large. Furthermore, when $|x| \to 1$, $\delta\phi(x)$ is sufficiently large. Thus, $|\mathcal{A}_0|$ is sufficiently close to 0. Therefore, $\mathcal{A}_0$ must be a set of volume zero, and $B^*$ fulfils the condition of Eq. (8).

**6. Prove the existence of diffeomorphic solutions in Eq. (17) and Eq. (18).**

We can obtain the same results for Eq. (17) and Eq. (18) based on the above proofs.

### SI-3. Data preprocessing and training details

#### SI-3.1. Simulation data

**Topup settings.** As the simulation data exclusively features susceptibility artifacts without within-subject head motion, we utilized the pre-defined configuration file (located at ${FSLDIR}/etc/flirtsch/b02b0.cnf) for running the Topup program, excluding head motion estimation.

**Network training settings.** We directly trained the network from scratch with a batch size of 1 over 3000 epochs at three scale levels (1000 epochs for each scale level). We set $\beta = 0.0001$, $\delta = 0.01$, $\gamma_1 = 0.0001$ and $\gamma_2 = 0.00005$ for all levels.

**Post-processing settings.** Since the simulation data doesn't include eddy current-induced distortions, any method that calculates the inhomogeneity field enables us to directly use the FSL FUGUE tool to remove susceptibility artifacts (SAs) in DWIs without the need for eddy current-induced distortion correction.

#### SI-3.2. HCP dataset

**Preprocessing.** We adopted the HCP pipelines (Glasser et al., 2013) to extract a single pair of b0 images distorted along the inverse-PE directions and preprocessed the T1w/T2w images. Then, the structural (T1w/T2w) images and the paired b0 images were rigidly co-registered.

**Network training settings.** The data of 300 of the 380 subjects were randomly selected as the training set, the data of 40 subjects were selected as the validation set, and the data of 40 subjects were selected as the testing set. We trained the network with a batch size of 5 over 450 epochs at three scale levels (150 epochs for each scale level). We set $\alpha = 1$, $\beta = 0.0001$, $\gamma_1 = 0.0001$ and $\delta = 0.01$ for all levels, set $\gamma_2 = 0.0001$ in the first two levels and set $\gamma_2 = 0.0005$ in the third level.

##### SI-3.3. HCP-D dataset

**Preprocessing.** We adopted the same preprocessing strategy as used for the HCP dataset to preprocess the HCP-D dataset.

**Network training settings.** The data of 544 of the 644 subjects were randomly selected as the training set, the data of 50 subjects were selected as the validation set, and the data of 50 subjects were selected as the testing set. We trained the network with a batch size of 5 over 400 epochs at three scale levels (150 epochs for the first two levels and 100 epochs for the last level). We set $\alpha = 1$, $\beta = 0.001$, $\delta = 0.01$ and $\gamma_1 = \gamma_2 = 0.0005$ for all levels.

##### SI-3.4. dHCP dataset

**Preprocessing.** We leveraged the dHCP minimal processing pipeline to extract b0 images distorted along 4 PE directions (LR, RL, AP, PA) (Bastiani et al., 2019) and preprocessed the T2w images (Makropoulos et al., 2018). The T2w and b0 images along the 4 PE directions were rigidly co-registered.

**Network training settings.** The data of 364 of the 444 subjects were randomly selected as the training set, the data of 40 subjects were selected as the validation set, and the data of 40 subjects were selected as the testing set. We independently trained and validated the model based on the AP-PA PE direction image sets and LR-RL PE direction image sets. For each pair of PE direction images, the network was trained with a batch size of 10 over 450 epochs at three scale levels (150 epochs for each scale level), and we set $\alpha = 1$, $\beta = 0.01$, $\gamma_1 = \gamma_2 = 0.001$, and $\delta = 0.01$.

##### SI-3.5. CBD dataset

**Preprocessing.** We adopted the same preprocessing strategy as that used for the HCP dataset to preprocess the CBD dataset.

**Network training settings.** The data of 242 subjects with CBDP were selected as the training set, the data of 40 subjects were selected as the validation set, and the data of 40 subjects were selected as the testing set. The CBDH dataset with 134 subjects was selected as an additional testing set to evaluate the model generalizability at other acquisition sites. Since the CBD data have only one

PE direction, we set $\alpha = \gamma_2 = 0$, $\beta = \gamma_1 = 1$, and $\delta = 1e4$. We did not multiply the Jacobian determinant of the inhomogeneity field in the DEW module. The proposed model was trained with a batch size of 5 over 600 epochs (200 epochs per scale level).

##### SI-3.6. Multicenter dataset

**Preprocessing.** We adopted the same preprocessing strategy as for the HCP dataset to preprocess the multicenter dataset.

**Network training settings.** We used network weights trained based on the HCP-D dataset as the initial weights for fine-tuning the model. Specifically, we selected the three subjects scanned at the first center as the training and validation sets and then predicted the inhomogeneity fields for the whole multicenter dataset after model training. We trained the network with a batch size of 1 over 300 epochs at three scale levels (100 epochs for each scale level). We set $\alpha$=1, $\beta$=0.005, $\delta$=0.01 and $\gamma_1$=$\gamma_2$=0.0005 for all levels. The details of the fine-tuning strategy are described in SI-6.

#### SI-4. Implementation of the deep-learning network

We used a convolutional neural network $f_\theta$ (shown in Fig. S1A), which consists of an encoder and decoder with skip connections (Ronneberger et al., 2015), to predict the inhomogeneity field $B$. The input images were first concatenated, and then the image intensity was normalized to between 0 and 1 using a co-normalization layer. The normalized images were then fed into a convolutional layer to extract the initial feature maps. These feature maps were then passed through a series of pre-activation residual layers in the encoder-decoder architecture, and the final convolutional layer mapped the transformed feature maps to a 1-channel inhomogeneity field.

The implementation details of the network are described as follows. For intensity normalization, an image-specific co-normalization layer was adapted according to the input image sets. If $\{I_1, I_2, I_a\}$ or $\{I_1, I_2\}$ was input into the network, the co-normalization layer used min–max normalization with a minimum value of $\min(\min(I_1), \min(I_2))$ and a maximum value of $\max(\max(I_1), \max(I_2))$ for $I_1$ and $I_2$ and a minimum value of $\min(I_a)$ and a maximum value of $\max(I_a)$ for $I_a$. For the image set $\{I, I_a\}$, the co-normalization layer used the typical min–max normalization method. We leveraged the residual blocks to form the encoding and decoding paths. Each residual block consisted of two consecutive convolutional layers with kernel sizes of 3, and each convolution was followed by a Leakey ReLU layer (slope set to 0.2), as shown in Fig. S1B. The left part is a regular residual block with an identity residual connection to maintain the resolution at the same scale level. The right part is a downsampling residual block to reduce the resolution among different scale levels, and the stride of the second convolutional layer was 2, with a $3 \times 3 \times 3$ convolution layer with a stride of 2 used to replace the identity residual connection.

#### SI-5. Evaluation metrics

##### SI-5.1. Fractional anisotropy-based mean squared difference (FA-MSD)

We calculated FA-MSD by computing the mean of the squared difference (MSD) between corresponding voxels inside the brain in FA images along inversed PE directions. In an ideal situation, there would be no difference between the FAs in inverse PE directions. A lower FA-MSD indicated better SAC performance. Notably, for the dHCP dataset, we calculated the FA-MSD metric between the FA image derived from the corrected AP-PA dMRI data and the FA image derived from the corrected LR-RL dMRI data, as shown in Fig. S3A.

##### SI-5.2. FA-based standard deviation (FA-STD)

We calculated FA-STD by computing the standard deviation (STD) between corresponding voxels inside the brain in FA images along the 4 PE directions (Wu et al., 2008). A lower FA-STD indicated better SAC performance, as shown in Fig. S3B. In this paper, only the dHCP dataset was used to calculate the FA-STD.

##### SI-5.3. FA-based structural similarity (FA-SS)

We calculated FA-SS by computing the mean value of the normalized cross correlation (NCC, window size=3) between the FA map and T1w images of all voxels inside the brain. An ideal FA map should present higher structural similarity with T1w images. A higher FA-SS(T1w) value indicates better alignment between the FA map and structural image.

##### SI-5.4. b0-based error (b0-Err) and FA-based error (FA-Err)

We computed b0-Err (FA-Err) by measuring the mean square error between the corrected b0 (FA) image and ground truth b0 (FA) image for all corresponding voxels within the brain. As we only have one simulation image, we reported the mean and standard deviation values of b0-Err (FA-Err) for voxels inside the brain in our paper.

##### SI-5.5. Number of folding voxels (NFV)

For the inverse-PE datasets (including HCP, HCP-D, dHCP and multicenter datasets), the NFV was calculated by counting the number of voxels that did not satisfy Eq. (8). Notably, for the dHCP dataset, we independently calculated the NFV values for both the AP-PA data and LR-RL data, which were denoted as NFV (AP-PA) and NFV (LR-RL), respectively. For the single-PE datasets, the NFV was calculated by counting the number of voxels that did not satisfy Eq. (S8):

$$\partial_v B(\boldsymbol{p}) > -1 \tag{S8}$$

##### SI-5.6. Coefficient of variation (CV) and intraclass correlation coefficient (ICC)

The formula for CV is expressed as follows:

$$CV(\boldsymbol{p}) = \frac{\sigma(\boldsymbol{p})}{\mu(\boldsymbol{p})} \times 100 \tag{S9}$$

where $\sigma(\boldsymbol{p})$ denotes the standard deviation of voxel $\boldsymbol{p}$ across all the centers, and $\mu(\boldsymbol{p})$ denotes the mean values of voxel $\boldsymbol{p}$ across all the centers. We calculated the ICC with a two-way mixed-effects model:

$$ICC = \frac{MS_R - MS_E}{MS_R + (k-1)MS_E + \frac{k}{n}(MS_C - MS_E)} \tag{S10}$$

where $MS_R$ denotes the mean square of between-subject variance, $MS_E$ denotes the mean square of error, $MS_C$ denotes the mean square of the within-subject variance, $n$ denotes the number of subjects, and $k$ denotes the number of centers per subject. In the case of the multicenter dataset, $n = 3$ and $k = 10$. Notably, we only calculated the CV and ICC values for the voxels inside the brain.

#### SI-6. The fine-tuning strategy

The fine-tuning strategy was similar to the training strategy. First, we loaded the pretrained model and initialized the network for each scale level with the pretrained weights. Then, we independently trained the network for each scale level. Once the current level's training was complete, we did not initialize the next level's network weights with the trained weights from the current level.

#### SI-7. Statistical results and additional figures

**Table S1. The quantitative results of b0-MSD and b0-SS on HCP dataset.**

| Dataset | Method | b0-MSD (1e6) ↓ | b0-SS ↑ |
|---|---|---|---|
| HCP | Topup | 0.521 (0.076) | 0.394 (0.040) |
| | S-Net | 2.502 (0.551) | 0.391 (0.027) |
| | SACNet(wos) | **0.072 (0.019)*** | 0.392 (0.027) |

| | | | |
|---|---|---|---|
| | SACNet(T1w) | 0.077 (0.020) | 0.404 (0.028) |
| | SACNet(T2w) | 0.092 (0.025) | **0.411 (0.032)*** |

Note: The median value is presented in the table, and the value in parenthesis represents the median absolute deviation. Bold font denotes that the method performs the best among the considered methods. The symbol "*" indicates that the method which achieved the best performance is significantly better than baseline methods (including Topup and S-Net), with p-value<0.05 (one-sided Wilcoxon signed-rank test). Statistical details between each two methods are provided in Table S3. "PED" denotes the phase encoding direction. " ↓ " denotes that the metric the lower the better, " ↑ " denotes that the metric the higher the better.

**Table S2. Statistical comparison of the Topup, S-Net, SACNet(wos) and SACNet(T2w) methods based on the simulated dMRI dataset.**

| | b0-Err | FA-Err | FM-Err |
|---|---|---|---|
| Pairwise comparison in severe distortion areas (voxels with distortions ≥10 mm) | p value (one-sided Wilcoxon rank test, right-sided) | p value (one-sided Wilcoxon rank test, right-sided) | p value (one-sided Wilcoxon rank test, right-sided) |
| SACNet(T2w) vs. SACNet(wos) | <0.001* | <0.001* | 1.000 |
| SACNet(T2w) vs. S-Net | <0.001* | <0.001* | <0.001* |
| SACNet(T2w) vs. Topup | 1.000 | 1.000 | 1.000 |
| SACNet(wos) vs. S-Net | <0.001* | <0.001* | <0.001* |
| SACNet(wos) vs. Topup | 1.000 | 1.000 | 1.000 |

**Table S3. Statistical comparison of the Topup, S-Net, SACNet(wos), SACNet(T1w) and SACNet(T2w) methods based on the HCP dataset.**

| | FA-MSD | FA-SS |
|---|---|---|
| Pairwise comparison | p value (one-sided Wilcoxon rank test, right-sided) | p value (one-sided Wilcoxon rank test, left-sided) |
| SACNet(T2w) vs. SACNet(T1w) | <0.001* | <0.001* |
| SACNet(T2w) vs. SACNet(wos) | <0.001* | <0.001* |
| SACNet(T2w) vs. S-Net | <0.001* | <0.001* |
| SACNet(T2w) vs. Topup | <0.001* | <0.001* |
| SACNet(T1w) vs. SACNet(wos) | <0.001* | <0.001* |

| | | |
|---|---|---|
| SACNet(T1w) vs. S-Net | <0.001* | <0.001* |
| SACNet(T1w) vs. Topup | <0.001* | <0.001* |
| SACNet(wos) vs. S-Net | <0.001* | <0.001* |
| SACNet(wos) vs. Topup | 0.027 | <0.001* |
| | **b0-MSD** | **b0-SS** |
| Pairwise comparison | p value (one-sided Wilcoxon rank test, right-sided) | p value (one-sided Wilcoxon rank test, left-sided) |
| SACNet(T2w) vs. SACNet(T1w) | 1.000 | <0.001* |
| SACNet(T2w) vs. SACNet(wos) | 1.000 | <0.001* |
| SACNet(T2w) vs. S-Net | <0.001* | <0.001* |
| SACNet(T2w) vs. Topup | <0.001* | <0.001* |
| SACNet(T1w) vs. SACNet(wos) | 1.000 | <0.001* |
| SACNet(T1w) vs. S-Net | <0.001* | <0.001* |
| SACNet(T1w) vs. Topup | <0.001* | 0.003 |
| SACNet(wos) vs. S-Net | <0.001* | 0.163 |
| SACNet(wos) vs. Topup | <0.001* | 0.574 |
| | **EPI-MSD** | **EPI-SS** |
| Pairwise comparison | p value (one-sided Wilcoxon rank test, right-sided) | p value (one-sided Wilcoxon rank test, left-sided) |
| SACNet(T2w) vs. SACNet(T1w) | <0.001* | <0.001* |
| SACNet(T2w) vs. SACNet(wos) | 1.000 | <0.001* |
| SACNet(T2w) vs. S-Net | <0.001* | <0.001* |
| SACNet(T2w) vs. Topup | <0.001* | 0.020 |
| SACNet(T1w) vs. SACNet(wos) | 1.000 | <0.001* |
| SACNet(T1w) vs. S-Net | <0.001* | <0.001* |
| SACNet(T1w) vs. Topup | 1.000 | 0.506 |
| SACNet(wos) vs. S-Net | <0.001* | <0.001* |
| SACNet(wos) vs. Topup | <0.001* | 0.839 |
| | **NFV (dMRI data)** | **NFV (Spin-echo Fieldmap)** |
| Pairwise comparison | p value (one-sided Wilcoxon rank test, right-sided) | p value (one-sided Wilcoxon rank test, left-sided) |
| SACNet(T2w) vs. SACNet(T1w) | 0.824 | 0.982 |
| SACNet(T2w) vs. SACNet(wos) | 0.712 | 0.660 |
| SACNet(T2w) vs. S-Net | <0.001* | <0.001* |

| | | |
|---|---|---|
| SACNet(T2w) vs. Topup | <0.001* | 0.004 |
| SACNet(T1w) vs. SACNet(wos) | 0.324 | 0.024 |
| SACNet(T1w) vs. S-Net | <0.001* | <0.001* |
| SACNet(T1w) vs. Topup | <0.001* | <0.001* |
| SACNet(wos) vs. S-Net | <0.001* | <0.001* |
| SACNet(wos) vs. Topup | <0.001* | 0.004 |

**Table S4. Statistical comparison of the Topup, S-Net, SACNet(wos), SACNet(T1w) and SACNet(T2w) methods based on the HCP-D dataset.**

| | **FA-MSD** | **FA-SS** | **NFV** |
|---|---|---|---|
| Pairwise comparison | p value (one-sided Wilcoxon rank test, right-sided) | p value (one-sided Wilcoxon rank test, left-sided) | p value (one-sided Wilcoxon rank test, right-sided) |
| SACNet(T2w) vs. SACNet(T1w) | <0.001* | 0.058 | 0.063 |
| SACNet(T2w) vs. SACNet(wos) | <0.001* | <0.001* | 1.000 |
| SACNet(T2w) vs. S-Net | <0.001* | <0.001* | <0.001* |
| SACNet(T2w) vs. Topup | <0.001* | <0.001* | <0.001* |
| SACNet(T1w) vs. SACNet(wos) | <0.001* | <0.001* | 1.000 |
| SACNet(T1w) vs. S-Net | <0.001* | <0.001* | <0.001* |
| SACNet(T1w) vs. Topup | <0.001* | <0.001* | <0.001* |
| SACNet(wos) vs. S-Net | <0.001* | <0.001* | <0.001* |
| SACNet(wos) vs. Topup | <0.001* | 0.005 | <0.001* |

**Table S5. Statistical comparison of the Topup, S-Net, SACNet(wos) and SACNet(T2w) methods based on the dHCP dataset.**

| | **FA-MSD** | **FA-STD** | **NFV (LR-RL)** | **NFV (AP-PA)** |
|---|---|---|---|---|
| Pairwise comparison | p value (one-sided Wilcoxon rank test, right-sided) | p value (one-sided Wilcoxon rank test, right-sided) | p value (one-sided Wilcoxon rank test, right-sided) | p value (one-sided Wilcoxon rank test, right-sided) |
| SACNet(T2w) vs. SACNet(wos) | <0.001* | <0.001* | 0.159 | 0.159 |

| | | | | |
|---|---|---|---|---|
| SACNet(T2w) vs. S-Net | <0.001* | <0.001* | <0.001* | <0.001* |
| SACNet(T2w) vs. Topup | <0.001* | <0.001* | <0.001* | <0.001* |
| SACNet(wos) vs. S-Net | <0.001* | <0.001* | <0.001* | <0.001* |
| SACNet(wos) vs. Topup | <0.001* | <0.001* | <0.001* | <0.001* |

**Table S6. Statistical comparison of the Fieldmap, VoxelMorph, SACNet(T1w) and SACNet(T2w) methods based on the CBD datasets.**

| | **FA-SS of CBDP** | **NFV of CBDP** | **FA-SS of CBDH** | **NFV of CBDH** |
|---|---|---|---|---|
| Pairwise comparison | p value (one-sided Wilcoxon rank test, left-sided) | p value (one-sided Wilcoxon rank test, right-sided) | p value (one-sided Wilcoxon rank test, left-sided) | p value (one-sided Wilcoxon rank test, right-sided) |
| SACNet(T2w) vs. SACNet(T1w) | 1.000 | N/A | 1.000 | N/A |
| SACNet(T2w) vs. VoxelMorph | <0.001* | <0.001* | <0.001* | <0.001* |
| SACNet(T2w) vs. Fieldmap | <0.001* | <0.001* | 0.005 | <0.001* |
| SACNet(T1w) vs. VoxelMorph | <0.001* | <0.001* | <0.001* | <0.001* |
| SACNet(T1w) vs. Fieldmap | <0.001* | <0.001* | <0.001* | <0.001* |

**Table S7. Statistical comparison of the Topup, S-Net, SACNet(wos) and SACNet(T1w) methods based on the multicenter dataset.**

| | **FA-MSD** | **FA-SS** | **NFV** |
|---|---|---|---|
| Pairwise comparison | p value (one-sided Wilcoxon rank test, right-sided) | p value (one-sided Wilcoxon rank test, left-sided) | p value (one-sided Wilcoxon rank test, right-sided) |
| SACNet(T1w) vs. SACNet(wos) | 0.026 | <0.001* | 0.988 |
| SACNet(T1w) vs. S-Net | <0.001* | <0.001* | <0.001* |
| SACNet(T1w) vs. Topup | <0.001* | 0.003 | <0.001* |

| | | | |
|---|---|---|---|
| SACNet(wos) vs. S-Net | <0.001* | <0.001* | <0.001* |
| SACNet(wos) vs. Topup | <0.001* | 0.031 | <0.001* |

**Table S8. Statistical comparison of the CV distributions of different subjects for the Topup, S-Net, SACNet(wos) and SACNet(T1w) methods based on the multicenter dataset.**

| | Subject-1 | Subject-2 | Subject-3 |
|---|---|---|---|
| Pairwise comparison p value (one-sided Wilcoxon rank test, right-sided) | p value | p value | p value |
| SACNet(T1w) vs. SACNet(wos) | 0.004 | <0.001* | <0.001* |
| SACNet(T1w) vs. S-Net | <0.001* | <0.001* | <0.001* |
| SACNet(T1w) vs. Topup | <0.001* | <0.001* | <0.001* |
| SACNet(wos) vs. S-Net | <0.001* | <0.001* | <0.001* |
| SACNet(wos) vs. Topup | <0.001* | <0.001* | <0.001* |

**Table S9. Statistical comparison of the ICC distributions in different brain regions for the Topup, S-Net, SACNet(wos) and SACNet(T1w) methods based on the multicenter dataset.**

| | Frontal | Parietal | Temporal | Occipital | Subcortical |
|---|---|---|---|---|---|
| Pairwise comparison (one-sided Wilcoxon rank test, left-sided) | p value | p value | p value | p value | p value |
| SACNet(T1w) vs. SACNet(wos) | <0.001* | 1.000 | <0.001* | 0.085 | 0.062 |
| SACNet(T1w) vs. S-Net | <0.001* | <0.001* | <0.001* | <0.001* | <0.001* |
| SACNet(T1w) vs. Topup | <0.001* | <0.001* | <0.001* | <0.001* | <0.001* |
| SACNet(wos) vs. S-Net | <0.001* | <0.001* | <0.001* | <0.001* | <0.001* |
| SACNet(wos) vs. Topup | <0.001* | <0.001* | <0.001* | <0.001* | <0.001* |

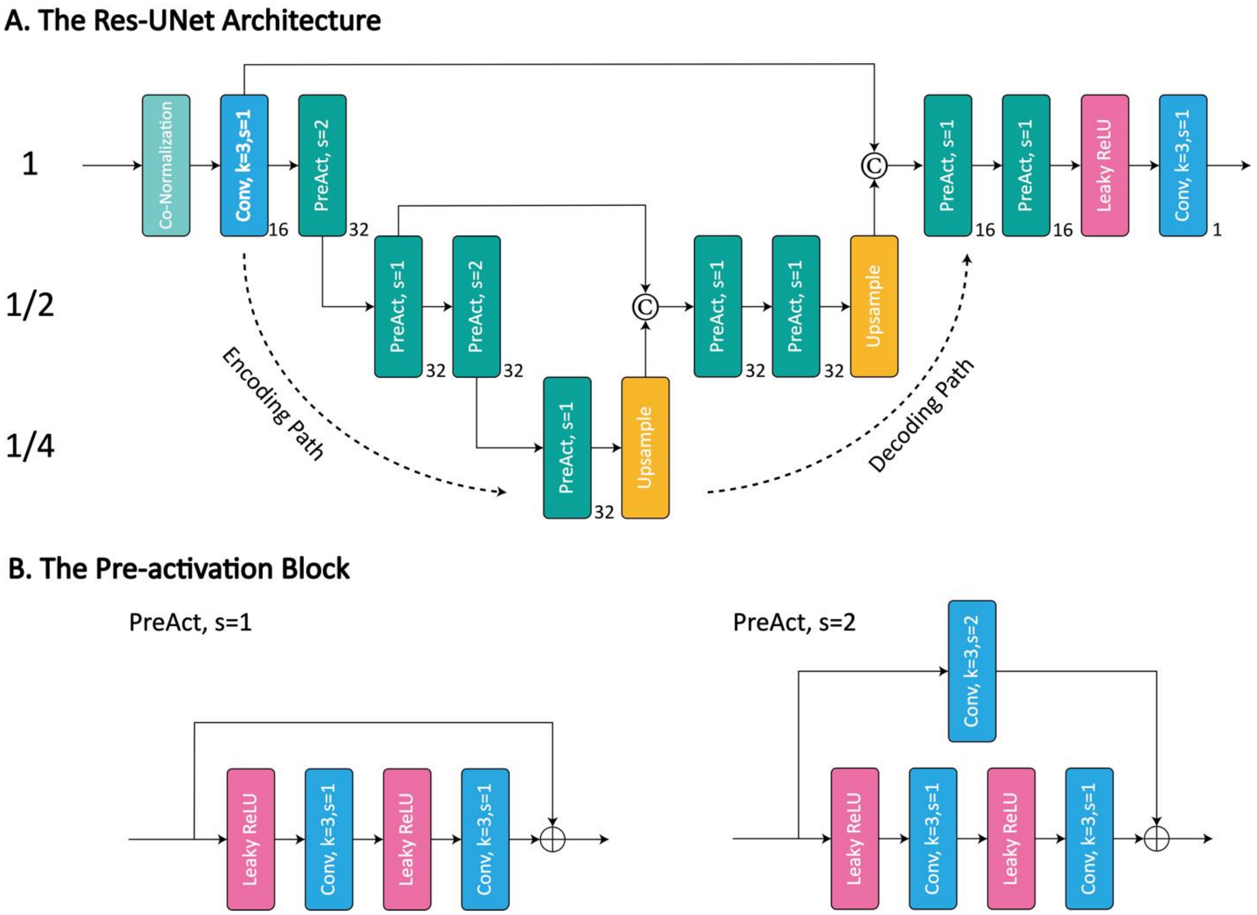


**Figure S1.** (A) Architecture of Res-UNet. The number in the lower right corner of each rounded rectangle represents the number of feature maps. (B) The pre-activation (PreAct) block used in the Res-UNet architecture.

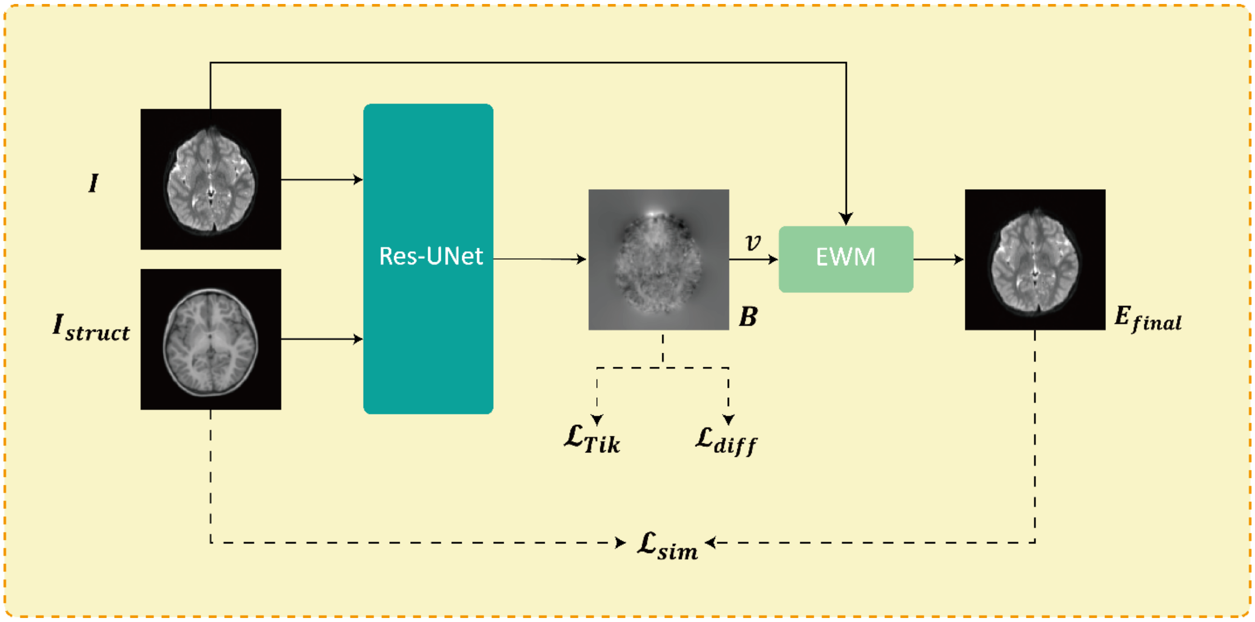


**Figure S2.** Diagram of the loss structure when only one b0 image and one structural image were used as inputs.

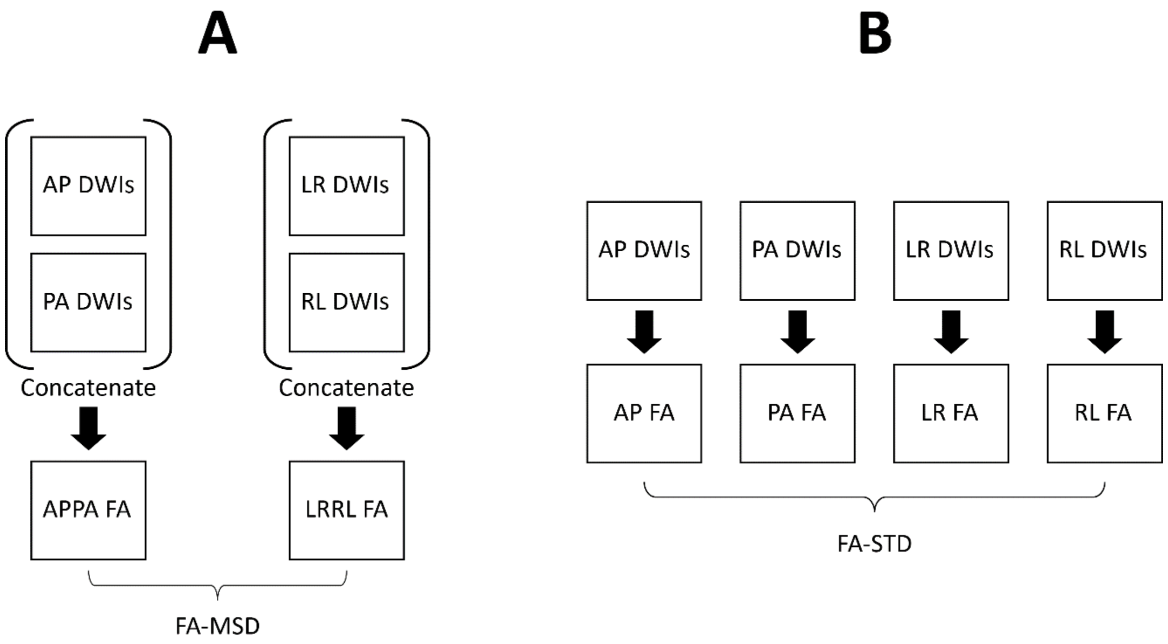


**Figure S3.** Illustration of the evaluation process for the dHCP dataset, including (A) the FA-MSD calculation process and (B) FA-STD calculation process.

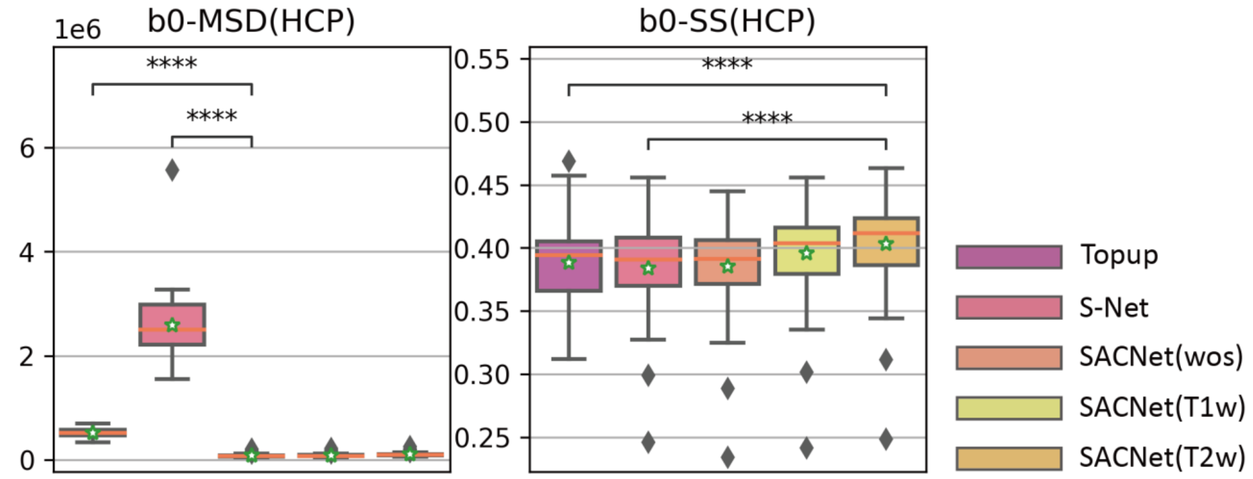


**Figure S4.** The quantitative results calculated on HCP b0 images. For all the boxplots, the green star indicates the mean value, and the coral horizontal line indicates the median value. We marked the significance of group differences between the best-performing model and baseline models. “****” denotes that $p \leqslant 0.0001$.

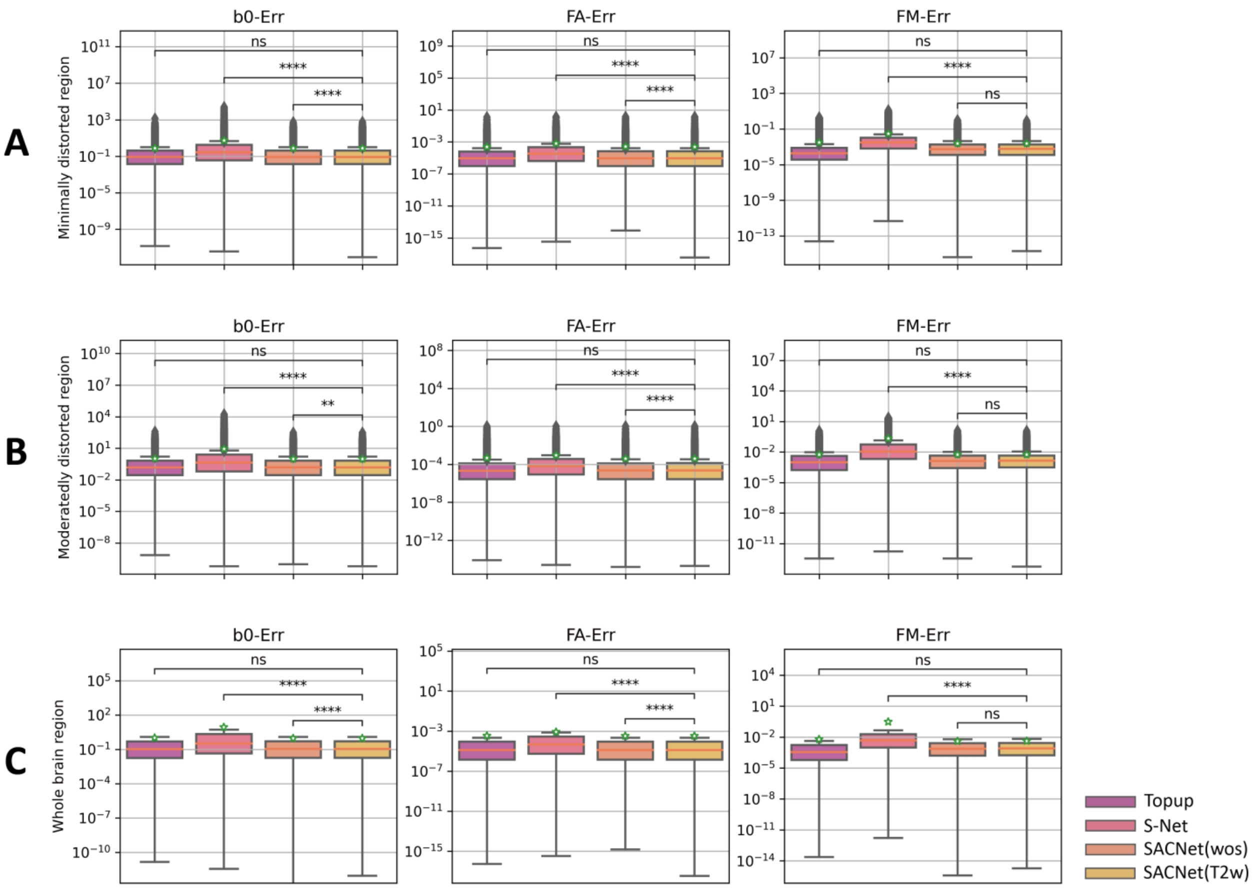


**Figure S5.** Boxplots showing quantitative results across different methods on brain regions with varying distortion levels on simulated dMRI images: (A) minimally distorted regions (voxels with distortions ≤ 2 mm), (B) moderately distorted regions (voxels with distortions between 2 mm and 10 mm), and (C) the entire brain region. FM represents fieldmap. The green star marks the mean value, while the coral horizontal line represents the median value. We marked the significance of group differences between the best-performing model and baseline models. “ns” denotes that $0.05 < p \leqslant 1$, “**” denotes that $0.001 < p \leqslant 0.01$, and “****” denotes that $p \leqslant 0.0001$.
